# Robust Self-Regeneratable Stiff Living Materials

**DOI:** 10.1101/2020.09.24.311589

**Authors:** Avinash Manjula-Basavanna, Anna Duraj-Thatte, Neel S. Joshi

## Abstract

Living systems have not only the exemplary capability to fabricate materials (*e.g.* wood, bone) under ambient conditions but they also consist of living cells that imbue them with properties like growth and self-regeneration. Like a seed that can grow into a sturdy living wood, we wondered: can living cells alone serve as the primary building block to fabricate stiff materials? Here we report the fabrication of stiff living materials (SLMs) produced entirely from microbial cells, without the incorporation of any structural biopolymers (*e.g.* cellulose, chitin, collagen) or biominerals (*e.g.* hydroxyapatite, calcium carbonate) that are known to impart stiffness to biological materials. Remarkably, SLMs are also lightweight, strong, resistant to organic solvents and can self-regenerate. This living materials technology can serve as a powerful biomanufacturing platform to design and develop sustainable structural materials, biosensors, self-regulators, self-healing and environment-responsive smart materials.

---

Innovation in materials science and technology has been a major driving force for the advancement of human civilization.(*1*) However, until relatively recently, materials innovations were pursued and implemented without much regard for global sustainability.(*2*) This must change rapidly in the face of the urgent threats of global warming and potentially irreversible ecological damage. Unlike our linear materials economy, which follows a make-use-dispose model, biology provides a template for a circular materials economy, in which abundant feedstocks are directed by cells into living materials that involve either regeneration or biodegradation at the end of the material’s life cycle. In most cases, these structure-building processes occur at ambient temperature and pressure, without the need for “heat-beat-treat” modalities of materials processing that are a hallmark of synthetic materials.(*3–6*) In many cases, biology accomplishes this task solely by using living cells and their byproducts as structural building blocks. The ability to adapt this approach to structure building for the purposes of scalable materials fabrication would be of great benefit to the development of more sustainable manufacturing practices.

In the last few decades, living cells have been engineered extensively to produce a wide variety of small molecules, polymers, drugs and fuels.(*7*) More recently, cells have also been engineered to produce functional materials directly and/or modulate their properties, which has led to the emergence of the new field of engineered living materials (ELMs).(*8–11*) Early examples of ELMs demonstrated binding to synthetic materials (*e.g.* stainless steel), templating nanoparticles (*e.g.* gold) and immobilizing enzymes (*e.g.* amylase).(*12, 13*) Additional work has focused on ELMs that function as catalytic surfaces, filtration membranes, under-water adhesives, pressure sensors, conductive films, gut adhesives, *etc*. (*14–27*) In spite of rapid progress in the field, it should be noted that there are few examples of ELMs that combine the structure-building capabilities of cells with their other capabilities, like self-regeneration, self-healing, or environmental responsiveness. ELM technologies that can streamline fabrication by relying more on autonomous cellular functions could help advance the field from a fundamental perspective and lead to ELMs compatible with scalable manufacturing techniques. Here we report the fabrication of a new class of ELMs wherein microbial cells serve as the sole structural building block. These materials are not only one of the stiffest known ELMs, but they can also self-regenerate, suggesting that they could fit into a model of a circular material economy (Fig. 1A).

**Fig. 1.**
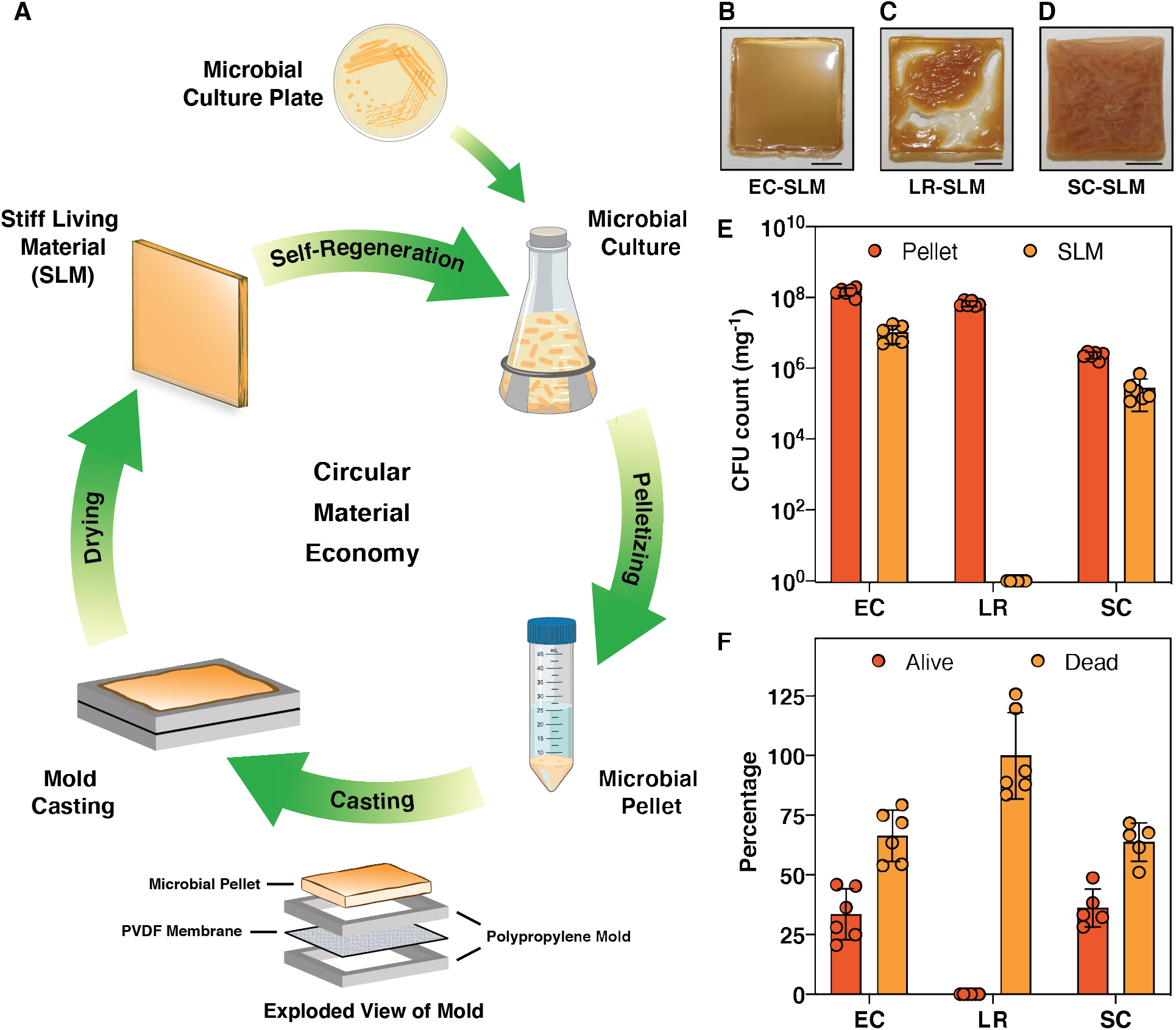
Fabrication of stiff living materials (SLMs). (A) Schematic shows the various stages involved in the fabrication and life cycle of SLMs produced solely from microbial cells. PVDF-Polyvinylidene fluoride. Optical images of SLM fabricated at 25 °C and 40±5 % relative humidity by air-drying for 24 h from (B) *Escherichia coli* (EC); (C) *Lactobacillus rhamnosus* (LR) and (D) *Saccharomyces cerevisiae* (SC). Scale bar 0.5 cm. (E) Colony forming unit (CFU) analysis of SLMs and their microbial pellet precursors. (F) Percentage of live and dead cells estimated from the SLMs with respect to their pellet (dry weight corrected). Bars represent mean values and the error bars are standard deviation.

## Evolution and fabrication of SLMs

Many published examples of ELMs take the form of soft materials in the form of biofilms, semi-solids or hydrogels. In these examples, cells were either combined with synthetic polymers or biopolymers to create composites, or they were designed/selected to produce a specific extra-cellular matrix (e.g. curli fibers, cellulose).(*12–27*) In our recent work, we have shown that this approach can be used to fabricate macroscopic stiff (2-4 GPa) thin films by exploiting the stiff structural characteristics of curli fibers.(*28*) In contrast to the above approaches, we wondered whether living cells alone, without any extracellular matrix, could serve as the primary building block in ELM fabrication and thereby incorporate aspects of self-regeneration. In order to explore this possibility, we started with a strain of *Escherichia coli* (PQN4), derived from an MC4100 lineage of laboratory strains, that is known not to produce any extracellular matrix components such as curli fibers, fimbriae, flagella or cellulose.(*12*) After culturing for 24 h in lysogeny broth, the *E. coli* cells were pelleted and washed with deionized water to remove the spent media. The resulting pellet, when drop-cast onto a glass slide and allowed to dry under ambient conditions, resulted in a brittle, transparent living material that fragmented during the drying process (Fig. S2A). However, upon seeing the potential for higher quality materials to be made with this basic approach, we iteratively refined the fabrication protocol in order to obtain increasingly robust prototypes (Fig. S2). We found that increasing the amount of wet biomass starting material and using a mold could slightly decrease the fragmentation of the SLM. Although this led to a material with a glossy top surface (Fig. S2F), the bottom surface (Fig. S2G) of the material had patches of cells that were not dried effectively, inhibiting the formation of a cohesive material. We reasoned that the non-porous glass substrate was inhibiting effective drying on the bottom surface of the SLM.

Applying a similar drop casting protocol on porous substrates like copper or stainless-steel mesh led to more uniformly dried materials with less fragmentation but left an imprint on the bottom surface of the SLM (Fig. S2K). Further attempts to use polymeric porous substrates either led to strong adhesion that prevented the removal of the SLM (e.g. nylon), or SLMs with curved architectures (e.g. PTFE-coated stainless steel), presumably due to different drying rates on the top and bottom surfaces. We then reasoned that a combination of vacuum suction and a substrate with balanced adhesion strength could lead to flat, cohesive materials that could be removed from the substrate and be self-standing. Accordingly, we applied the drying protocol to a polyvinylidene fluoride (PVDF) membrane that is typically used to bind proteins in western blots. By drop casting the *E. coli* cell pellet on a PVDF membrane mounted on a Millipore SNAP i.d. Mini Blot Holder connected to low vacuum suction, we were able to achieve a flat, mostly cohesive material, though some fragments still formed (Fig. S2U,V). Counterintuitively, the same setup in the absence of suction led to fragmentation-free, flat SLMs. Although the adhesion of the SLM to the PVDF membrane prevented it from being removed manually, we found that the membrane could be removed by applying a small amount of dimethylformamide (DMF), which is known to solubilize PVDF. We also tried using higher temperatures (50/75/100 °C) to speed up SLM formation, but this led to samples with extensive cracks and discoloration or charring (Fig. S2N-T).

An optimized fabrication of the SLM involved firmly sandwiching the PVDF membrane between two polypropylene molds, then casting the *E. coli* cell pellet on top of the membrane and drying under ambient conditions (25 °C and 40±5 % relative humidity) for 24 h. This yielded a fragmentation-free glossy flat SLM of 0.4-1.2 mm thickness (Fig. 1B). Given that the precursor material to the SLM consisted solely of live *E. coli* cells (EC-SLM), we were curious about how the fabrication protocol affected cell viability. Remarkably, one milligram of EC-SLM was found to have 1.0+0.5 ~10^7^ colony forming units (CFUs), while its precursor (i.e. the wet cell pellet) had a CFU count of 1.5+0.04 *10^8^ mg^-1^ (Fig. 1E). We then employed the same protocols to the Gram-positive bacterium *Lactobacillus rhamnosus* and the yeast *Saccharomyces cerevisiae* to investigate whether other microbes can also form SLMs in a similar manner. Interestingly, *L. rhamnosus* resulted in a SLM (LR-SLM) with a wrinkled top surface, while the SLM from *S. cerevisiae* (SC-SLM) had extensive cracks and a non-glossy texture (Fig. 1C,D). CFU analysis revealed that SC-SLM had 2.7+0.2 *10^5^ mg^-1^, but no cells were found to be alive in the LR-SLM (Fig. 1E). Since a large fraction of the wet weight of the cell pellets was water, we normalized the CFU counts for all three SLMs to the dry mass of the cell pellets (Fig. S6,7) and found that 33.5%, 0% and 36.1% of the original cells were alive in EC-SLM, LR-SLM and SC-SLM, respectively (Fig. 1F). Thus, we have demonstrated the very first examples of living bulk materials fabricated entirely from viable microbial cells. Further, in order to probe whether biomass from lysed cells could produce materials with similar qualitative properties, we used *E. coli* cells that had been lysed by exposure to 70% ethanol. After drying, these materials were not able to maintain a cohesive structure and as expected no living cells were recoverable by CFU analysis (Fig. S8).

## Physical characteristics of SLMs

Given the highly heterogeneous molecular composition of SLMs, we sought to understand their structure with a range of analytical techniques. X-ray diffraction (XRD) analysis of SLMs indicated that both EC-SLM and LR-SLM have a main diffraction peak corresponding to a *d*-spacing value of 0.44 nm, while EC-SLM has two additional peaks at 0.88 nm and 0.23 nm (Fig. 2A). Although it is difficult to assign the identity of these peaks, the spectra do establish that SLMs are amorphous materials. Thermal gravimetric analysis (TGA) showed a slightly negative slope below 100 °C that is likely due to water loss, followed by degradation above 130 °C (Fig. S9). Differential scanning calorimetry (DSC) analysis of EC-SLM showed a glass-transition-like second-order transition (50-60 °C) during the first cycle of the heating curve (Fig. S10). However, the successive heat-cool cycles did not reveal the presence of such transitions, suggesting the probable role of water acting as a plasticizer. Similar features were also observed in DSC traces of LR-SLM and SC-SLM (Fig. S11). EC-SLM appeared to be somewhat transparent by qualitative observation, but the absorption spectrum showed less than 10% transparency, across the visible range (Fig. S13). Based on the cell viability experiments showing that only ~35% of the cells in EC-SLM were alive after the fabrication protocol, we reasoned that the remaining cells likely lysed during the drying process, releasing their periplasmic and cytosolic contents. This mixture, containing nucleic acids, lipids, and proteins assembles into an amorphous network, with residual water serving as a plasticizer.

**Fig. 2.**
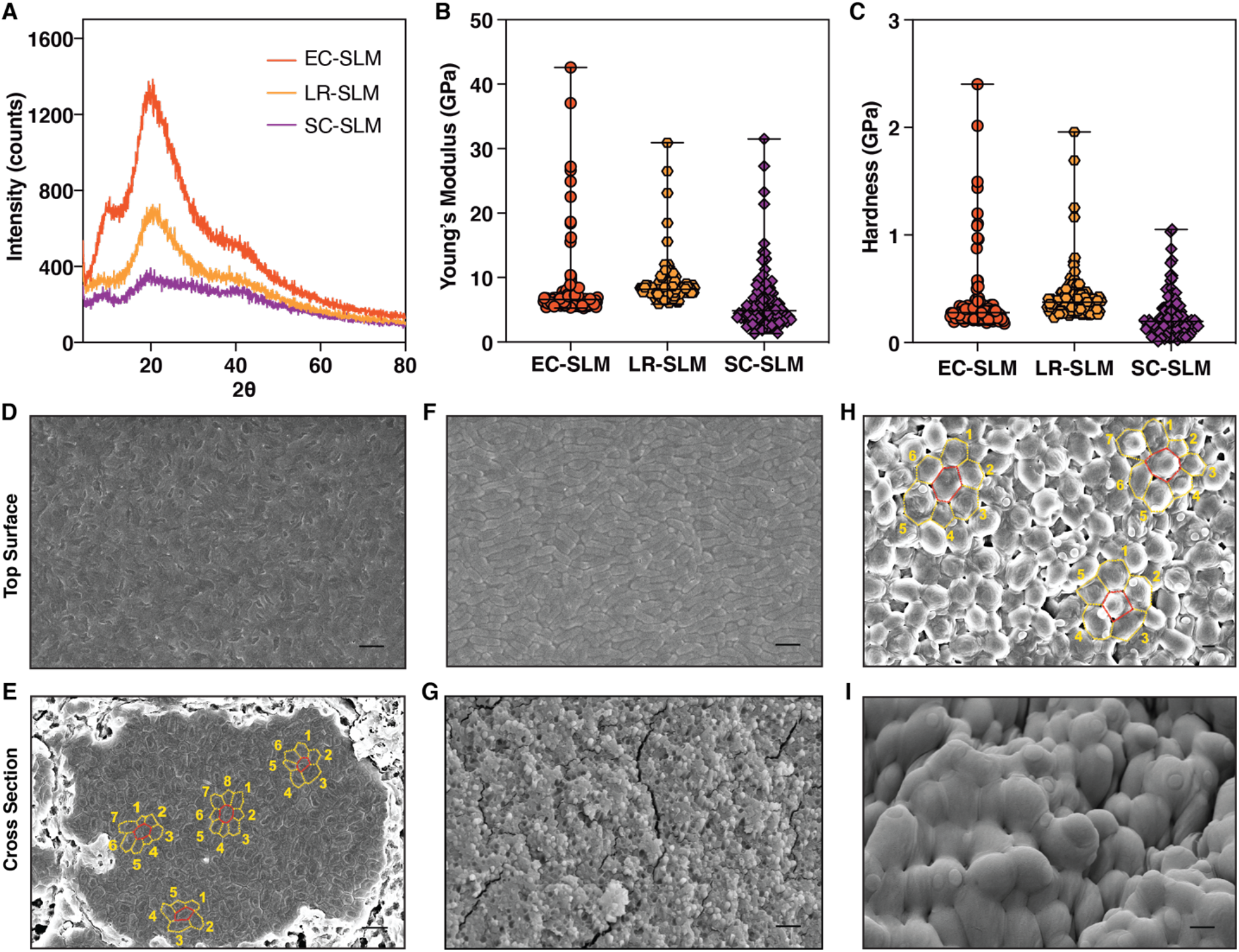
Physical and structural characteristics of SLMs. (A) X-ray diffraction spectra of SLMs of various microbial composition. (B) Young’s modulus and (C) Hardness of SLMs obtained from nanoindentation (n≥125). The graphs show median and the range. Field emission scanning electron microscopy (FESEM) images of EC-SLM (D,E); LR-SLM (F,G) and SC-SLM (H,I). (D,F,H) Top surface of SLM. (E,G,I) Cross-section of SLM. Scale bar 2 μm. (E,H) show the planar packing density, *η* (number of neighboring cells within the same plane).

## Mechanical characteristics of SLMs

The mechanical properties of the SLMs were investigated by nanoindentation, as it uses small loads that are suitable for biomaterials and can probe heterogeneity in microscopic dimensions.(*29, 30*) SLMs were indented (n≥125) with a Berkovich diamond tip to obtain the continuous load, *P*, versus depth of penetration, *h*, curves. Nanoindentation experiments showed smooth *P-h* curves, which were analyzed using the Oliver-Pharr method to extract Young’s modulus, *E*, and hardness, *H*, of the SLMs (Fig. S14). EC-SLM was found to have *E* ranging from 5 to 42 GPa, while their *H* were about 0.2 to 2.4 GPa (Fig. 2B,C). LR-SLM and SC-SLM also showed stiffness and hardness values in a similar range as EC-SLM (Fig. 2B,C). The mechanical characteristics of SLMs were consistent across different samples, while the slightly wider distribution of stiffness could be attributed to the heterogeneous components and their packing (Fig. S15). Interestingly, the SLM obtained from lysed *E. coli* (70% ethanol treatment) also exhibited similar *E* and *H* values, which further supports that cellular components can self-assemble, albeit heterogeneously, to form stiff materials (Fig. S16).

## Morphological characteristics of SLMs

In order to understand how the structure of SLM materials, formed exclusively from microbial cells, was able to remain stiff while preserving cell viability, we turned to electron microscopy. Field emission scanning electron microscopy (FESEM) imaging of the top surface of EC-SLM revealed a dense matrix of *E. coli* cells which all appear to be ruptured (Fig. 2D). But from CFU analysis, we know that *E. coli* cells are alive in EC-SLM, prompting us to investigate the core of the material. Cross-sectional imaging of EC-SLM showed a fascinating ordering of cells into tightly packed domains amidst loosely bound cells (Fig. 2E). Each domain can comprise of anywhere between 3 to nearly 500 cells, spanning up to a width of 30 μm. It should be noted that *E. coli* is a rod-shaped cell but, in these domains, transforms to a polygonal prism with a planar packing density (*η*, number of neighboring cells within the same plane) of predominantly 6, although 5, 7 and 8 are also observed (Fig. 2E). We speculate that the cells in the loosely bound regions have greater survivability compared to the tightly packed domains. Contrastingly, the top surface of LR-SLM was found to have an array of *L. rhamnosus* cells, whose rod-shaped structure appeared to be intact (Fig. 2F). It is difficult to ascertain the *η* of *L. rhamnosus* due to their known inherent tendency to form chains. The cross-sectional images of LR-SLM revealed that the cells were lysed to form an amorphous heterogenous solid (Fig. 2G). These FESEM images of LR-SLM provide additional evidence for the lack of CFU in the samples (Fig. 1E,F). In contrast, SC-SLM was found to form a close packing of spherical shaped *S. cerevisiae* cells with *η* of 5, 6 and 7 (Fig. 2H). Interestingly, the cross section of SC-SLM showed that *S. cerevisiae* cells were packed less densely at the core but formed tightly compressed layers both on the top and bottom surfaces (Fig. 2I, S19). Thus, it appears that lysis of *S. cerevisiae* cells forms a hard-protective shell on the outer surface and thereby enables cells at the core to survive to a greater extent.

## Self-regeneration of SLMs

We then exploited the living cells embedded in the SLMs to develop a protocol for self-regeneration. When a small fragment (5-10 mg) of EC-SLM was introduced into selective media, the SLM started to disperse and the embedded cells started proliferating to form a turbid culture. After 24 h of culture, the cells were pelleted and cast onto the mold as per the same fabrication protocol described above in order to create the second generation (Gen II) EC-SLM fabricated from a fragment of the first generation (Gen I) EC-SLM (Fig. 3A). The process could be repeated again to form a Gen III EC-SLM. Both Gen II and Gen III EC-SLMs were found to have a CFU count of around 10^7^ mg^-1^, which is almost same as that of Gen I (Fig. 3B). Moreover, nanoindentation studies showed that *E* (5-41 GPa) and *H* (0.2-2.5 GPa) of the Gen II and Gen III EC-SLMs were also similar to that of the Gen I sample (Fig. 3C,D). CFU analysis of EC-SLM samples stored on the benchtop over time revealed that the value decreased to ~10^4^ mg^-1^ on day 15 and 21 mg^-1^ on day 30 (Fig. 3E). We calculated an exponential cell death rate of 0.43 per day.

**Fig. 3.**
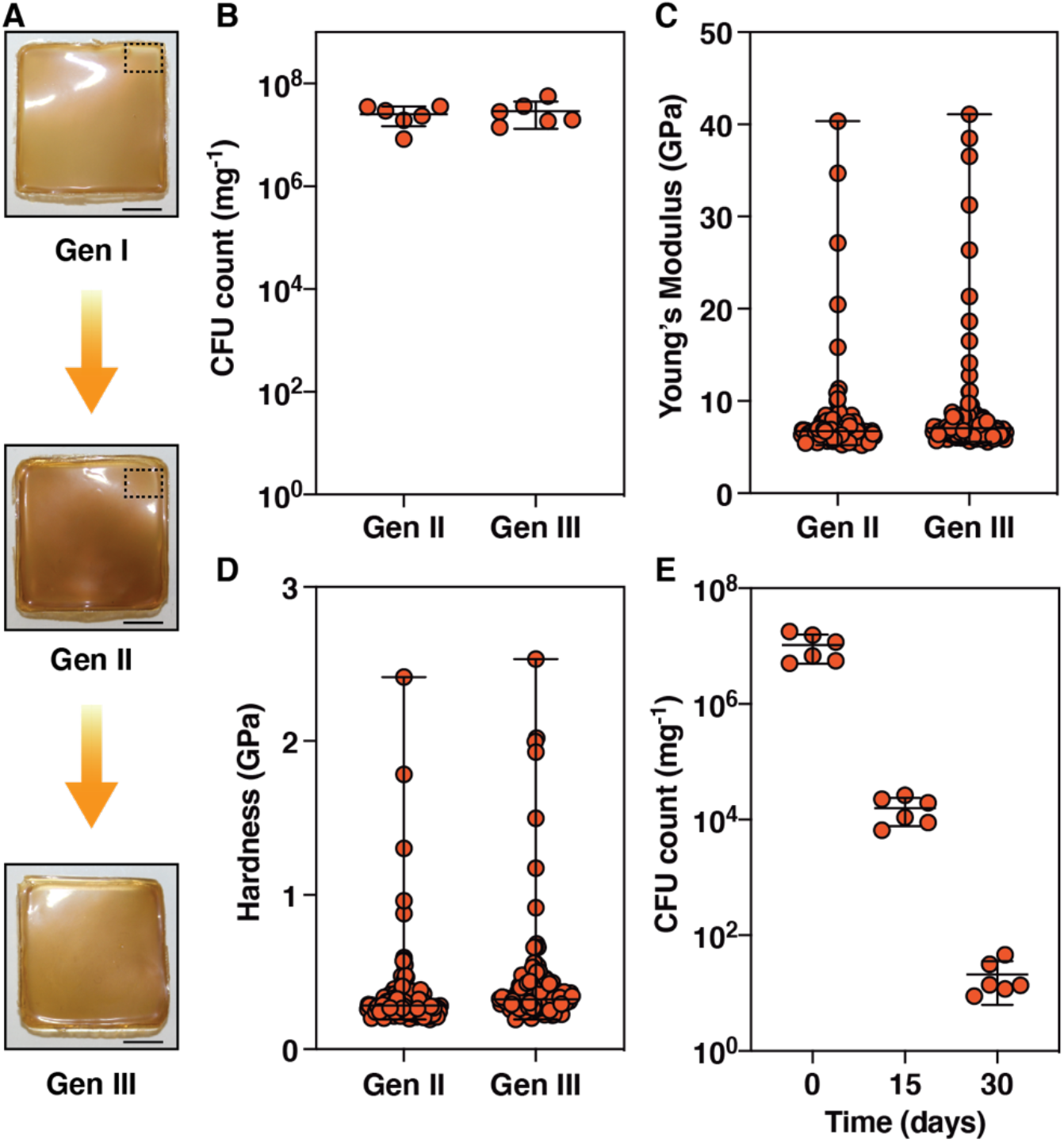
Self-regeneration of EC-SLM. (A) Optical images of first (Gen I), second (Gen II) and third (Gen III) generations of EC-SLM. A small fragment (dotted rectangle) of Gen I was cultured, pelletized and air-dried to produce the Gen II, which in turn resulted in Gen III. (B) CFU count of Gen II and Gen III of EC-SLM. (C) Young’s modulus and (D) Hardness of Gen II and Gen III EC-SLMs measured by nanoindentation (n≥125). The graphs show median and the range. (E) Time dependent CFU analysis of EC-SLM. The graph shows mean values and the error bars are standard deviation.

## Robustness of SLMs

In the process of optimizing our SLM fabrication protocol, we observed that the SLM does not appear to be affected by the DMF used to remove the PVDF membrane. We also noticed that EC-SLM did not disperse even when submerged in DMF. Intrigued by this observation, we submerged EC-SLM in a range of solvents with varying properties - hexane, chloroform, ethyl acetate, acetonitrile, absolute ethanol, methanol, DMF and deionized water (Fig. S20). EC-SLM dispersed only in water and was completely stable in all of the other solvents, whose polarity index spans the entire spectrum. After 24 h of submersion in the various solvents, we subjected the samples to CFU analysis and were surprised to find values >10^6^ mg^-1^ in all solvents except chloroform and methanol, which led to complete cell death (Fig. 4A). When we repeated the solvent submersion and CFU analysis with the wet *E. coli* cell pellet, we found that the cells died completely in all the solvents, except for hexane and deionized water (Fig. S21). The result with hexane may perhaps be explained by its complete lack of miscibility with water and lower density, allowing it to rest on top of the cell pellet. In contrast, EC-SLM was stable in both water-miscible and - immiscible organic solvents (Fig. 4B) and showed no significant weight loss (Fig. 4C) after 24 hours of incubation in various solvents, suggesting that the densely packed surface layer of the SLM, consisting of lysed cells, exhibits a protective effect on the cells embedded in the core. We also tested the robustness of EC-SLM by incubating it at 100 °C for 1 h and found a mean CFU count of over 700 mg^-1^ (Fig. 4D).

**Fig. 4.**
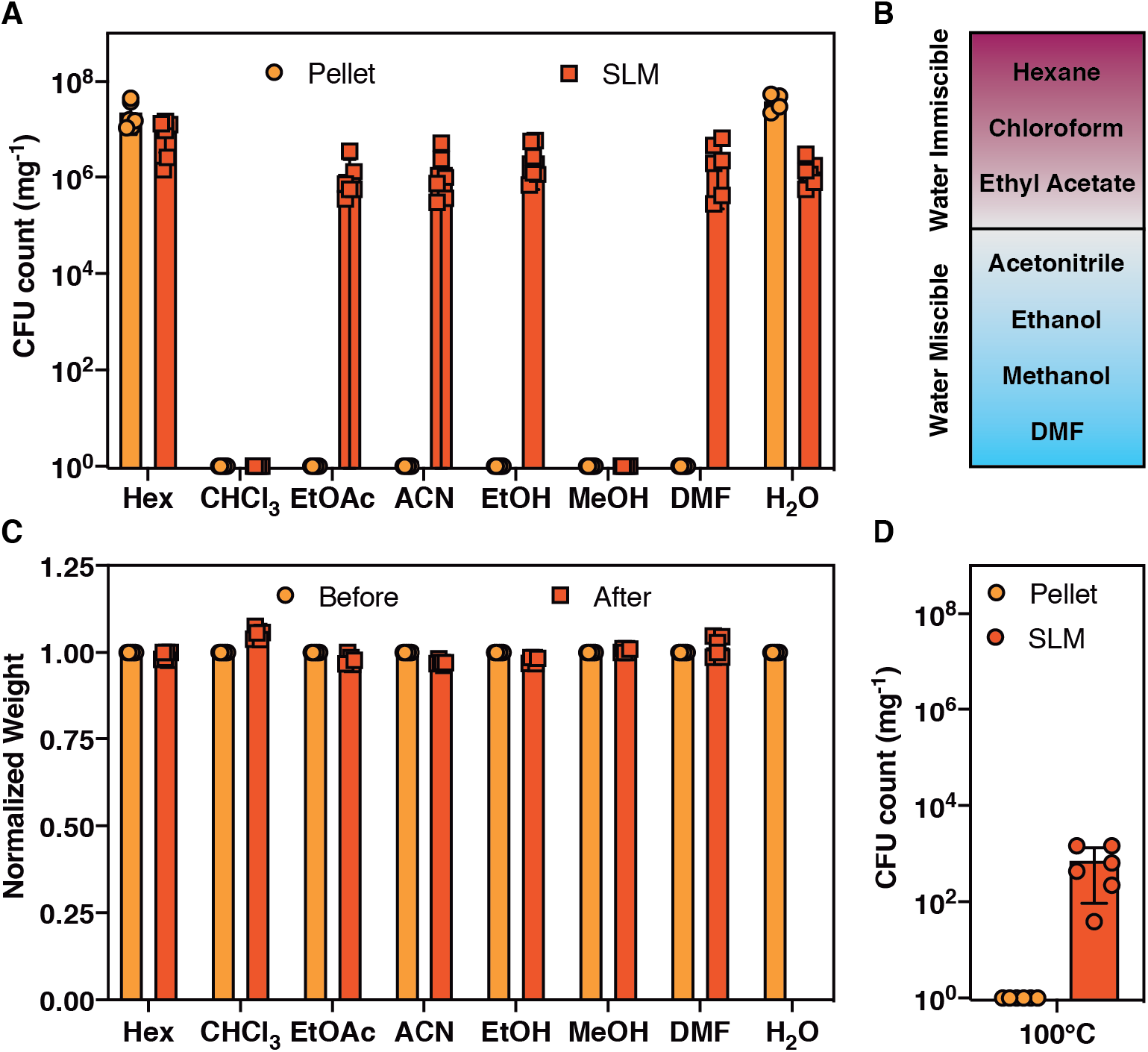
Solvent resistance of EC-SLM. (A) CFU count of EC-SLM and *E. coli* pellet after 24 h submersion in various solvents. CFU count of pellets were corrected for their dry weights. Hex: hexane; CHCl3: chloroform; EtOAc: ethyl acetate; ACN: acetonitrile, EtOH: absolute ethanol; MeOH: methanol; DMF: dimethylformamide. (B) Chart shows miscibility and immiscibility of organic solvents in water. (C) Normalized weights of EC-SLMs before and after 24 h submersion in solvents. (D) CFU count of EC-SLM and *E. coli* pellet after incubation at 100 °C for 1 h. The bar graphs represent mean values and the error bars are standard deviation.

## Mechanical landscape of SLMs

Based on our nanoindentation studies, it is evident that SLMs are remarkably stiff and hard for a material composed purely of microbial biomass. To put these properties into perspective, we provide a comparison of the mechanical properties of SLMs to other biomaterials and various types of human-made materials – metals, polymers, composites, ceramics, elastomers and foams. An Ashby plot of *E* versus density (*ρ*) shows that SLMs are stiffer than most biomaterials and polymers, and more comparable to composites (Fig. 5A).(*31*) We also obtained the yield strength, σy (estimated using the relation σy = *H*/3) of SLMs, which was found to be about 60-800 MPa.(*32, 33*) In Figure 5B, we show the Ashby plot of specific modulus (*E*/*ρ*) and specific strength (*σ*_y_/*ρ*), which indicates that the specific properties of SLMs are comparable to metals and ceramics, due to their low density.(*31*) Further, we provide specific examples of materials that are categorized into biomaterials (*e.g.* silk, collagen, cellulose *etc*.), biomaterials with cells (e.g. wood, skin, ligament *etc*.), non-biological materials (*e.g.* steel, glass, concrete, plastics *etc*.) and SLMs in an Ashby plot of *E* and strength, σ, in Figure 5C.(*31, 34*) Notably, the stiffness and strength of SLMs are comparable or superior to actin, balsa, cancellous bone, skin and plastics, amongst others, and they are comparable to robust structural materials such as silk, collagen, wood and concrete.

**Fig. 5.**
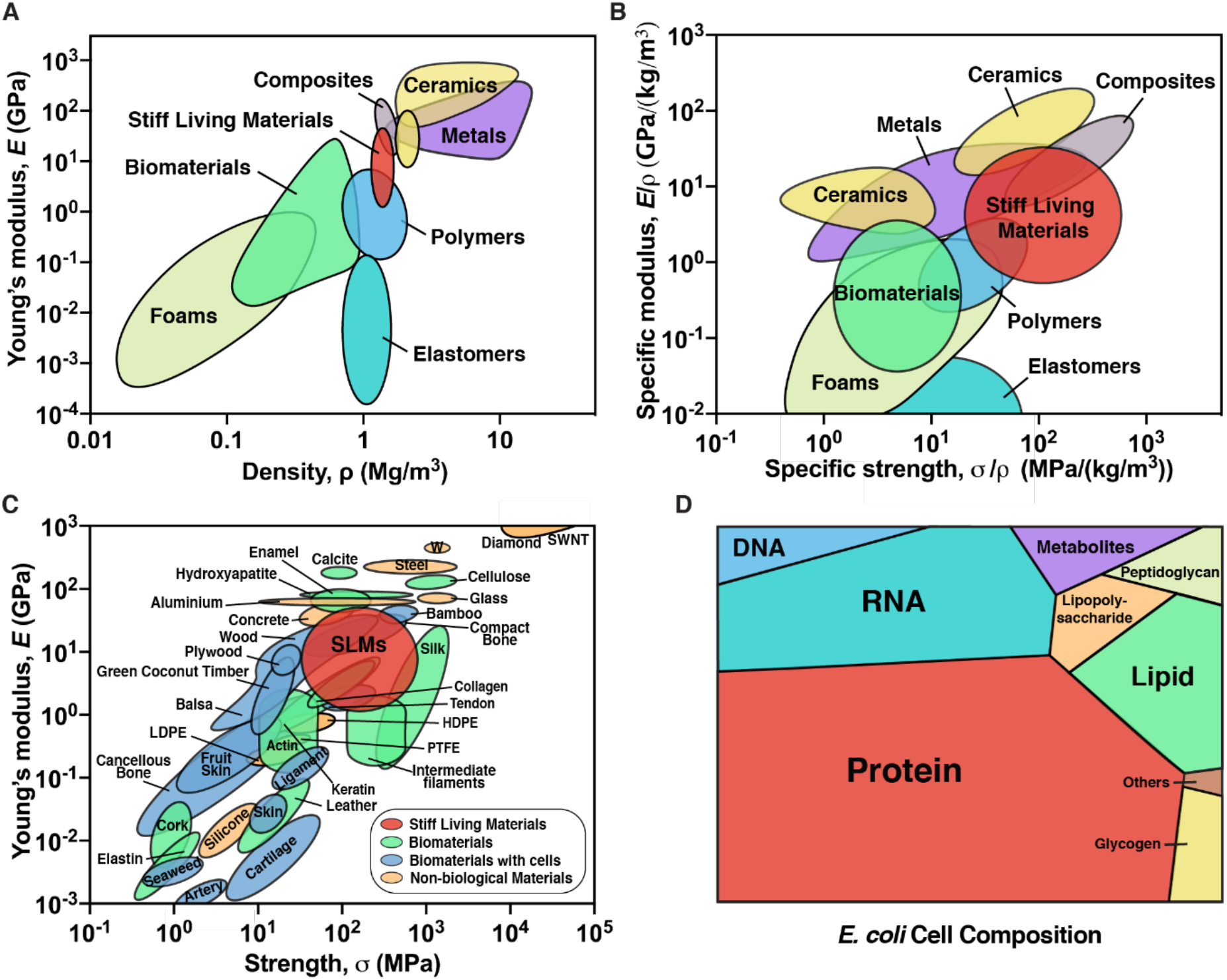
Mechanical and compositional landscape of SLMs. Ashby plot of (A) Young’s modulus verses density, (B) Specific modulus verses specific strength and (C) Young’s modulus verses strength, for various classes of materials and SLMs. SWNT: Single-wall carbon nanotube; LDPE: low density polyethylene; HDPE: high density polyethylene, PTFE: polytetrafluoroethylene. Adapted from references 33 and 34. (D) Voronoi tree diagram shows the relative amounts of the components present in the dry *E. coli* cell. Adapted from reference 35.

## Discussion

A living cell is a heterogenous mixture of proteins, nucleic acids, sugars *etc*. and to comprehend their contributions to SLM composition, we have presented a Voronoi tree diagram that shows the relative abundance of various components of a dry *E. coli* cell (Fig. 5D, Table S2).(*35*) Although it is difficult to ascertain the role of each of these cellular components in the formation of SLMs, the ability to do so from three cell types that vary widely in their anatomy and composition suggests that some of the observed properties of SLMs (e.g. stiffness, hardness, cohesiveness) may arise from a mixture of many different biomolecules. Other examples of stiff structural materials, like wood and bone, are composed of cells embedded in extracellular matrix components (e.g. cellulose, lignin, collagen, hydroxyapatite) with precise molecular compositions and self-assembly mechanisms that have been optimized over millions of years of evolution to exhibit mechanical robustness and other functional material properties. Given this, it is noteworthy that even non-specific mixtures of biomolecules derived from microbial cells can create rigid materials that rival their naturally occurring counterparts in terms of stiffness and hardness. The inability of ethanol treated *E. coli* cells (wherein ethanol disrupts the lipid membrane and denatures the proteins) to form a cohesive fragmentation-free SLM highlights that cellular integrity is essential and also suggests that active cellular processes that may play an important role in the formation of SLMs.

Indeed, many microbes have developed molecular, structural, metabolic and physiological adaptations to keep them alive under low-water conditions (i.e. xerotolerance).(*36–38*) Some of these xerotolerance mechanisms (*e.g.* production of trehalose, extracellular polymeric substances, hydrophilins *etc.*) may contribute to SLM material properties, and will be investigated in future studies. Although dried microbes have been widely used (*e.g.* dry baker’s yeast) for a very long time and mechanisms of microbial xerotolerance have been studied for decades, these studies were usually carried out in small volumes (*e.g.* microliters of microbial culture) and focused on either deciphering the mechanisms of xerotolerance or enhancing the survivability of microbes.(*38, 39*) Other research on microbial desiccation has focused on maximizing their long-term viability during storage, for example when probiotics are combined with emulsifiers and other additives prior to lyophilization.(*40*) In spite of all the fundamental knowledge and technological advancements around microbial drying processes, to the best of our knowledge, dried microbes have not been investigated as building blocks for macroscopic structural materials, as we demonstrate here.

Creating materials that incorporate microbes is a growing sub-field within synthetic biology and biomanufacturing. Most previous endeavors have accomplished this by either combining cells with exogenously supplied polymeric scaffolds or by engineering the cells to produce specific ECM components. Here we simplify this process to the extreme by using only living cells as a precursor to the material of interest and employ only drying under ambient conditions to achieve rigid, cohesive material. We have shown SLM fabrication to be compatible with several industrially relevant workhorse microbes, which could position it as a powerful biomanufacturing strategy that overcomes the inherent inefficiencies involving the separation of cells from their products. Given that the cells used in this work were not engineered in any way, the potential for using synthetic biology techniques to further enhance and tailor material properties is genuinely exciting. The incorporation of renewable feedstocks to fuel microbial growth could also position this approach as a promising manufacturing paradigm that is in line with a circular materials economy model.

## Acknowledgments

Authors thank Prof. Ju Li and Dr. Antoni Sanchez-Ferrer for helpful discussions. National Science Foundation 2026 Idea machine (Big Ideas) for stimulation and support. Work was performed in part at the Center for Nanoscale Systems at Harvard.

## Author contributions

A.M-B. conceived the project, designed, and performed all experiments. A.M.D-T. contributed to SLM fabrication, CFU and solvent resistance studies. All authors discussed and analyzed data. A.M-B. and N.S.J. wrote the manuscript.

## Data and materials availability

All data are available in the main text or the supplementary materials.

## Supplementary Materials

Materials and Methods

Figures S1-S21

Table S1 & S2

