## Supplemental Materials for "Robust Self-Regeneratable Stiff Living Materials"

**Contents**

- Materials and Methods
- Figures S1 to S21
- Table S1 & S2

### Materials and Methods

Cell Strains and Plasmids

SLMs were fabricated from the following cell strains;

- PQN4, *Escherichia coli* cell strain derived from LSR10 (MC4100, Δ*csgA*, λ(DE3), $Cam^R$). pET-21d(+) plasmid (Novagen 67743-3) was transformed to PQN4 to confer ampicillin resistance.
- *Lactobacillus rhamnosus* (ATCC® 27773™)
- *Saccharomyces cerevisiae* (ATCC® 9763™)

Fabrication of SLMs

*E. coli* (lysogeny broth with 100 μg/ml carbenicillin, 37 °C, 24 h), *L. rhamnosus* (MRS broth with 25 μg/ml chloramphenicol, 37 °C, 48 h) and *S. cerevisiae* (YPD broth, 30 °C, 24 h) were cultured (500 ml media) in a shaking incubator. *E. coli* and *S. cerevisiae* cells were pelletized at 3000 rpm, whereas 8000 rpm was employed for *L. rhamnosus*. The microbial cells were then washed twice (250 ml and 50 ml) with deionized water to remove the culture media. The resulting microbial pellet was casted on a PVDF (polyvinylidene fluoride; Millipore Immobilon-P, IPVH09120) membrane, which was firmly sandwiched between polypropylene molds (inner dimensions 2 cm X 2 cm X 1.6 mm) by adhesive tape. The microbial pellet was air-dried at ambient conditions (25 °C & 40±5 % relative humidity) for 24 h. The PVDF membrane strongly adhered to EC-SLM and LR-SLM, but did not stick to SC-SLM. By gently wiping the PVDF membrane with DMF (dimethylformamide) and waiting 5-10 min, it can be easily peeled off. The SLMs were stored at ambient conditions on the benchtop and used as needed.

Colony Forming Unit (CFU) Analysis of SLMs

5-10 mg of SLM or 20-100 mg of the microbial wet pellet (water washed) was subjected to serial dilutions and each dilution was plated onto a selective agar plate. The resulting colonies were counted to obtain the CFU count. For time-dependent CFU analysis, the SLMs stored at ambient conditions were utilized at day 0, 15 and 30.

Thermal Gravimetric Analysis (TGA)

TGA experiments were performed using a TA Q5000 IR instrument. SLMs (5-10 mg) were run at 5 °C $min^{-1}$ under $N_2$ purging at 50 mL $min^{-1}$ in platinum pans.

Dynamic Scanning Calorimetry (DSC)

DSC measurements were done using a TA Q200 instrument. Measurements were run under $N_2$ purging at 40 mL $min^{-1}$ and at 2 °C $min^{-1}$ with ~5 mg of SLM. Each measurement was performed in aluminum pans in the range of -10 to 100 °C with successive heat-cool cycles.

UV-Vis Absorption Spectroscopy

UV−Vis spectra were recorded on a Cary 5000 UV−Vis−NIR spectrophotometer (Agilent Technologies) in the range of 400 to 800 nm to obtain their percentage transparency.

X-Ray Diffraction (XRD):

XRD experiments on SLMs were performed using a Bruker D2 Phaser equipped with a beam of $\lambda_{CuK\alpha}$ = 0.15418 nm. The diffraction intensity of SLMs were recorded for $2\theta$ in the range of 4° to 80°.

Nanoindentation

Nanoindentation studies were performed on the samples using the Agilent Technologies G200 Nanoindenter. The instrument continuously monitors the load, $P$, and the depth of the penetration, $h$, of the indenter with the resolutions of 1 nN and 0.2 nm, respectively. A Berkovich diamond tip indenter with the tip radius of ~100 nm is used for the indentation. A peak load, $P_{max}$ of 1 mN with the loading and unloading rates of 0.2 mN $s^{-1}$, poisson's ratio of 0.3 and a hold time (at $P_{max}$) of 10 s was employed. A minimum of 125 indentations was performed in each case. The $P$-$h$ curves were analyzed using the Oliver-Pharr method to extract the Young's modulus ($E$), and the hardness ($H$) of the samples. The yield strength, $\sigma_y$ was estimated using the relation $\sigma_y = H/3$.

Field Emission Scanning Electron Microscope (FESEM)

FESEM samples were prepared by sputtering a 10-20 nm layer of Pt/Pd/Au. Images were acquired using a Zeiss Ultra55/Supra55VP FESEM equipped with a field emission gun operating at 5-10 kV.

Solvent Resistance of SLMs

EC-SLMs (~10 mg) were fully immersed in 2 ml of hexane, chloroform, ethyl acetate, acetonitrile, absolute ethanol, methanol, dimethylformamide (DMF) or deionized water. After 24 h of immersion, the SLMs were removed and air-dried overnight to remove any traces of solvent. The weight of EC-SLMs before and after the solvent immersion was noted before the samples were subjected to CFU analysis. As EC-SLM disperses in water, we could not obtain its weight after the submersion.

Self-regeneration of SLMs

5-10 mg of EC-SLM (first generation, Gen I) was added to 500 ml of lysogeny broth supplemented with 100 μg/ml carbenicillin incubated at 37 °C for 24 h. The cells were pelletized, casted on to the mold and air-dried to obtain the second generation, Gen II of EC-SLM (same fabrication protocol as described above). Similarly, a 5-10 mg fragment of Gen II was utilized to obtain the third generation, Gen III of EC-SLM.

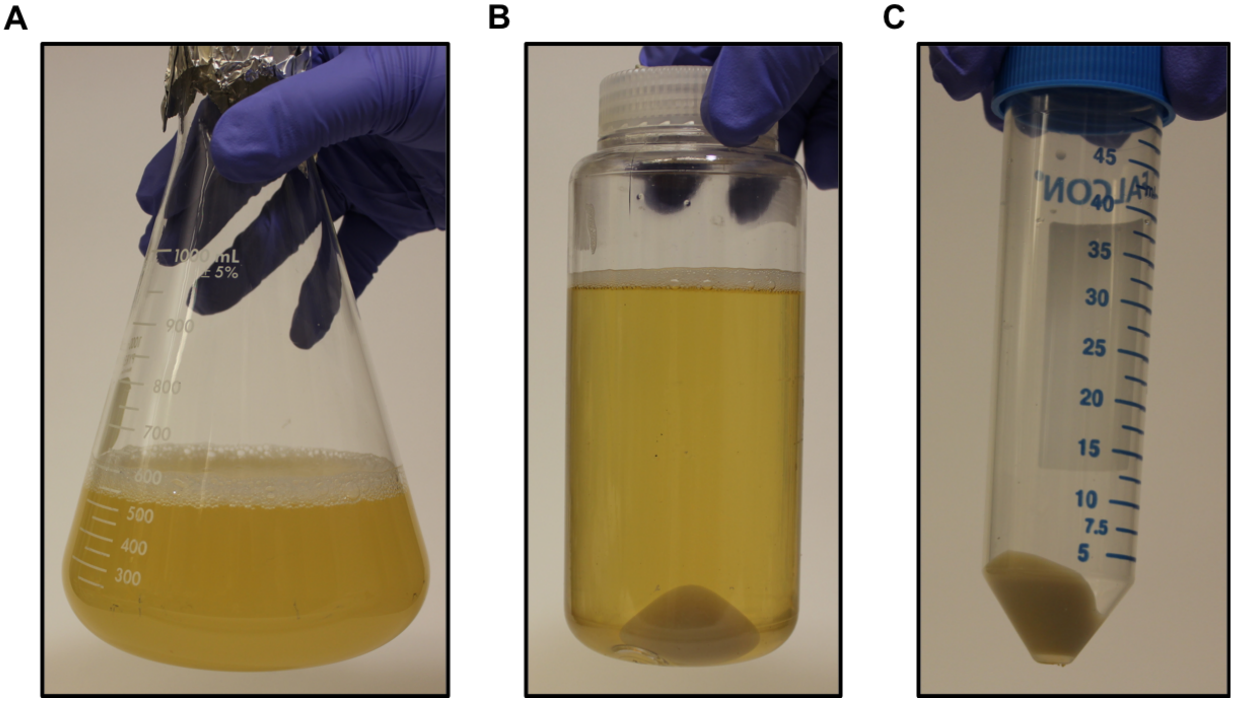


**Fig. S1. Optical images of microbial culture and pellet.** (A) *E. coli* culture in lysogeny broth after 24 h. (B) *E. coli* cells pelletized from the culture. (C) *E. coli* pellet after washing with deionized water.

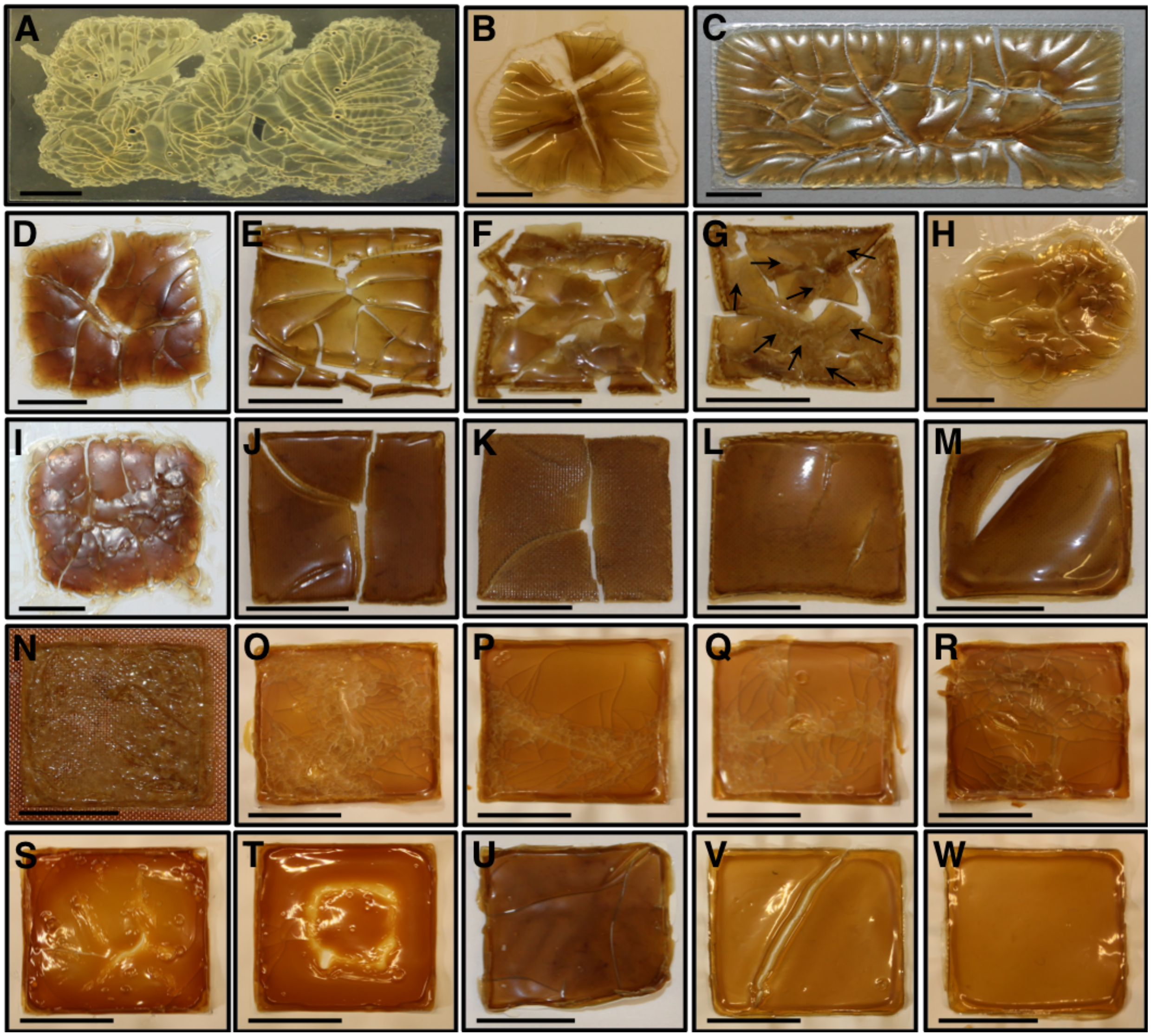


**Fig. S2. Optical images showing the evolution of stiff living material (SLM).**
(A-I) *E. coli* pellet dried on a glass substrate. (C,E-G) Pellet casted within a mold. (D-I) Higher pellet amount. (G) Arrows indicate ineffective drying of cells that inhibits formation of a cohesive glossy material. (H,I) Under low vacuum suction. (J,K) *E. coli* pellet dried on copper mesh substrate. *E. coli* pellet dried on stainless-steel mesh (L) and polytetrafluoroethylene (PTFE) coated steel mesh substrates (M). (N) *E. coli* pellet dried on copper mesh at 50 °C for 24h. (O-W) *E. coli* pellet dried on polyvinylidene fluoride (PVDF). Pellet dried at 50 °C for 24h (O), 75 °C for 2h (P), 75 °C for 3h (Q), 75 °C for 6h (R), 100 °C for 1h (S) and 100 °C for 2h (T). (U,V) Low vacuum suction using Millipore SNAP i.d. Mini Blot Holder. (W) Pellet dried at ambient conditions (25 °C, 24 h, 40±5 % humidity). (A-F,H-J,L-W) Top surface. (G,K) Bottom surface.

| Parameter | Types/Conditions | Remarks |
|---|---|---|
| Substrate | Glass | The bottom surface of SLM had scale-like architectures of patches of cells possibly due to ineffective drying on the non-porous nature of glass surface. SLM cracks extensively. |
| | Copper Mesh | The bottom surface of SLM lacked scale-like architectures. SLM cracks and it is non-flat. The mesh pattern gets imprinted on the bottom surface of SLM. |
| | Stainless Steel Mesh | The bottom surface of SLM lacked scale-like architectures. SLM cracks and it is non-flat. The mesh pattern gets imprinted on the bottom surface of SLM. |
| | Nylon Membrane | Adheres to nylon and the bottom surface of SLM had tiny scale-like architectures when peeled off from nylon. SLM Cracks. The mesh pattern gets imprinted on the bottom surface of SLM. |
| | Polytetrafluoroethylene (PTFE) Coated Steel Mesh | SLM does not stick to PTFE. SLM Cracks and it is deformed to a non-flat (curvy) shape. |
| | Polyvinylidene fluoride (PVDF) | SLM adheres to PVDF that cannot be peeled off manually but can be removed by gently wiping with dimethylformamide (DMF) solvent. Flat and fragmentation-free SLMs can be obtained. |
| Temperature | 25/50/75/100 °C | Higher temperature speeds up the SLM formation but have disadvantages like extensive cracks, charring/discoloration (depending on temperature and duration) and enhanced cell death. At 25 °C, the SLM formation takes up to 24 h. |
| Low Vacuum Suction | Millipore SNAP i.d. Mini Blot Holder<br><br>Vacuum Desiccator | SLM formation speeds up when vacuum suction (unidirectional; from bottom side of the cell pellet) is applied via a blot holder. Blot holder set up can facilitate flat SLMs. The three-dimensional suction in a vacuum desiccator results in potholes on SLM that cracks and it is non-flat (highly curvy). Vacuum desiccator takes longer time for SLM formation. |
| Duration | 12/24/48 h (at 25 °C)<br>1/2/3/6/24 h (at 50/75/100 °C) | 24 h duration was found to be optimal for SLM formation at 25 °C, while at 100 °C, 1-3 h was sufficient but suffer from cracking, charring and/or higher cell deaths. |
| Lysing Cell | 70% Ethanol Treatment<br>Ultrasonication<br>Freeze-Thawing | Ethanol treatment was found to be more convenient and effective than the other methods. |

**Table S1. List of parameters and conditions employed during the fabrication of SLM.**

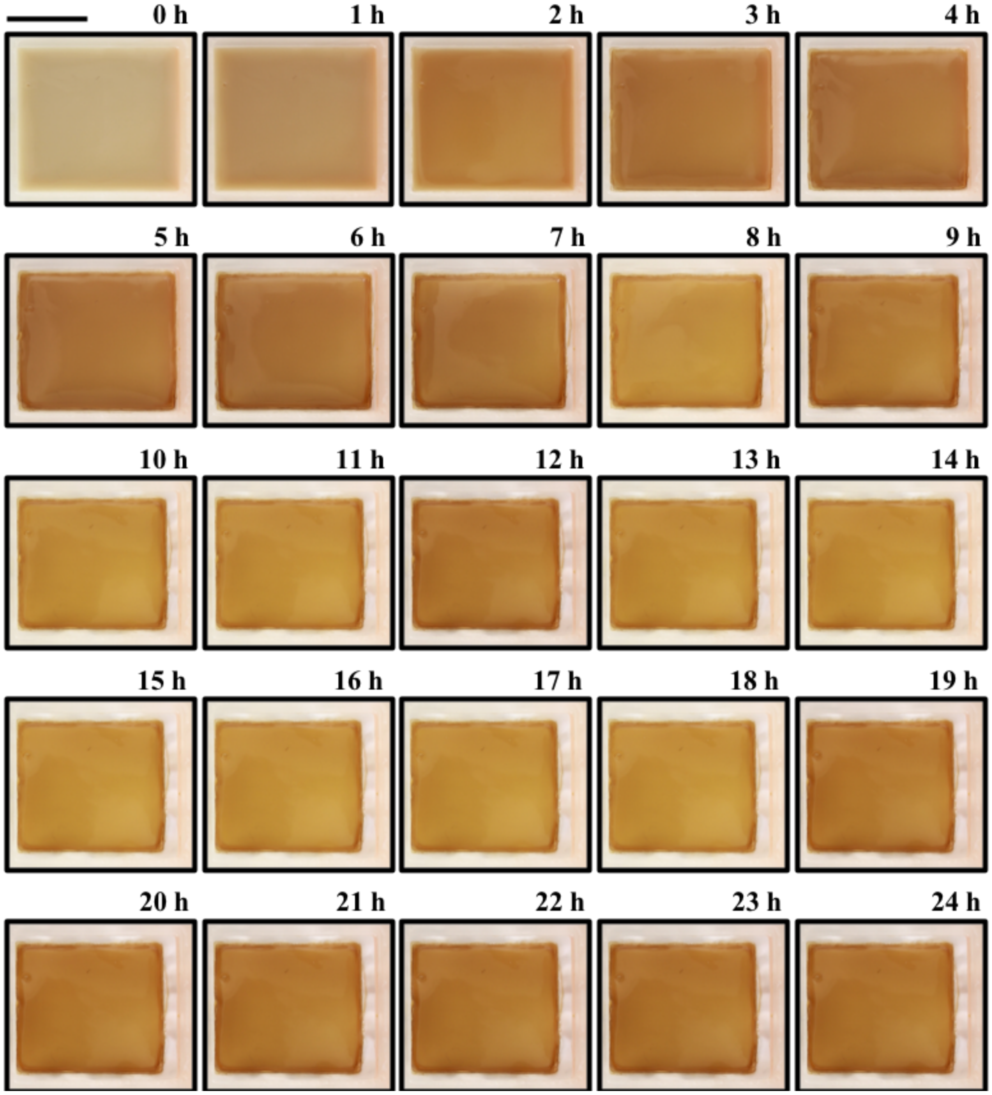


**Fig. S3. Optical time lapse images (recorded every hour) showing the ambient drying of EC-SLM.** Scale bar 1 cm.

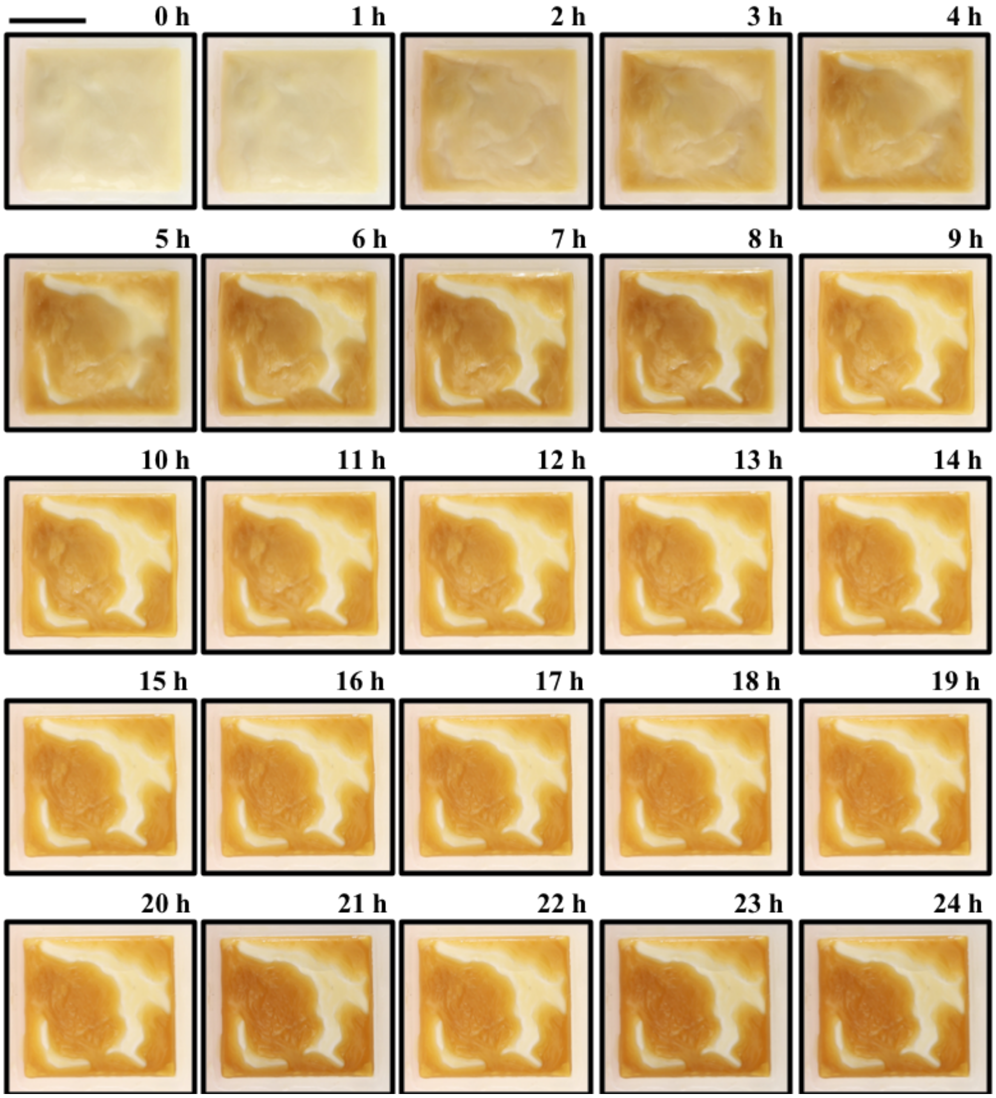


**Fig. S4. Optical time lapse images (recorded every hour) showing the ambient drying of LR-SLM.** Scale bar 1 cm.

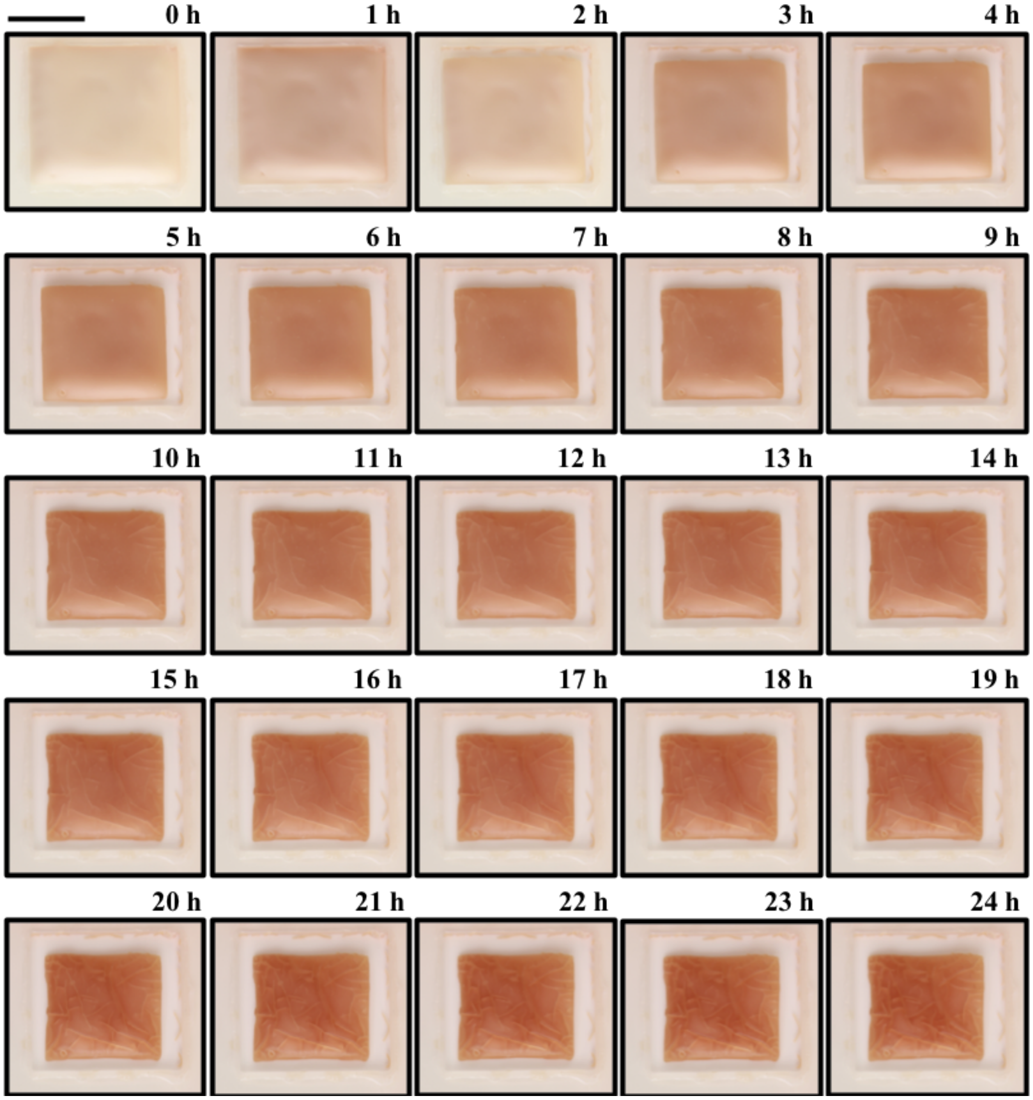


**Fig. S5. Optical time lapse images (recorded every hour) showing the ambient drying of SC-SLM.** Scale bar 1 cm.

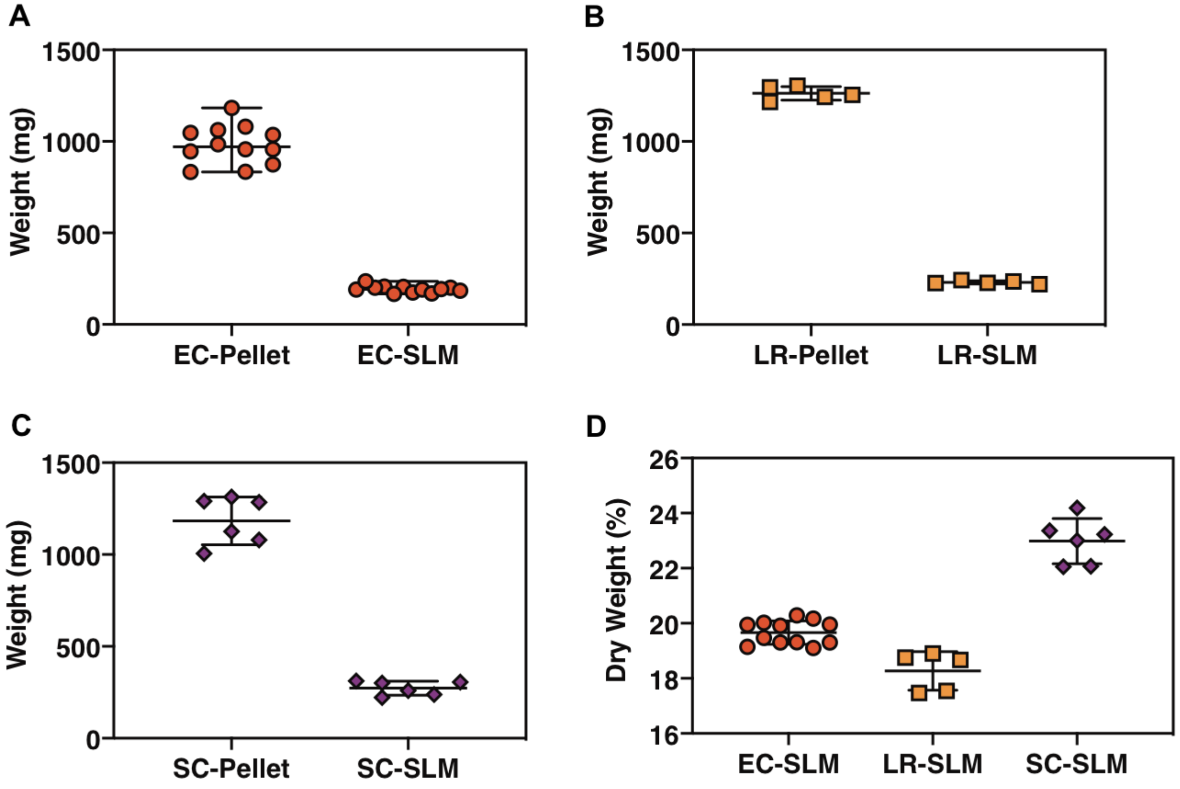


**Fig. S6. Weight analysis of SLM.** (A) *E. coli* pellet weight before and after drying (24 h) to form the SLM. (B) *L. rhamnosus* pellet weight before and after drying (24 h) to form the SLM. (C) *S. cerevisiae* pellet weight before and after drying (24 h) to form the SLM. (D) Dry weight percentage of EC-SLM, LR-SLM and SC-SLM. The graphs show mean values and the error bars are standard deviation.

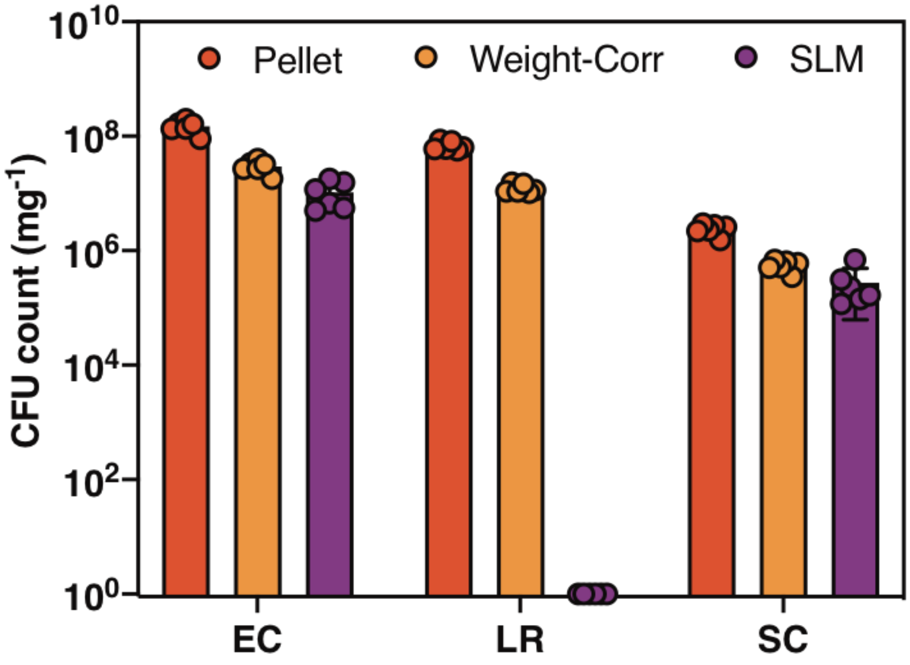


**Fig. S7. Colony forming unit (CFU) analysis of SLM.** CFU counts of *E. coli*, *L. rhamnosus* and *S. cerevisiae* of pellet, weight-corr (pellet corrected for dry weight as per Fig. S6D) and SLM. The bar graphs represent mean values and the error bars are standard deviation.

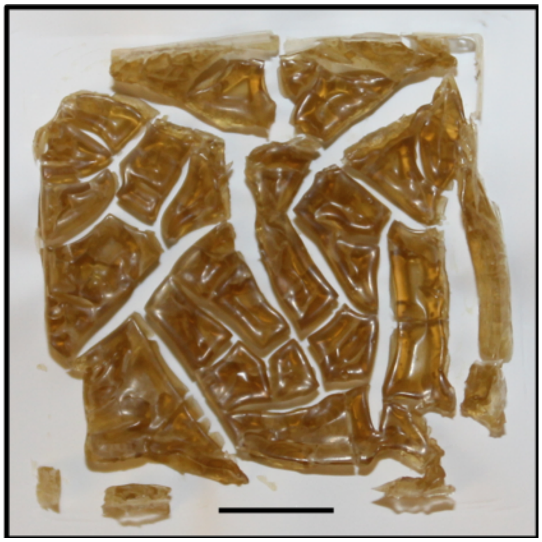

**Fig. S8. Optical image of SLM fabricated from 70% ethanol treated *E. coli* cells.**

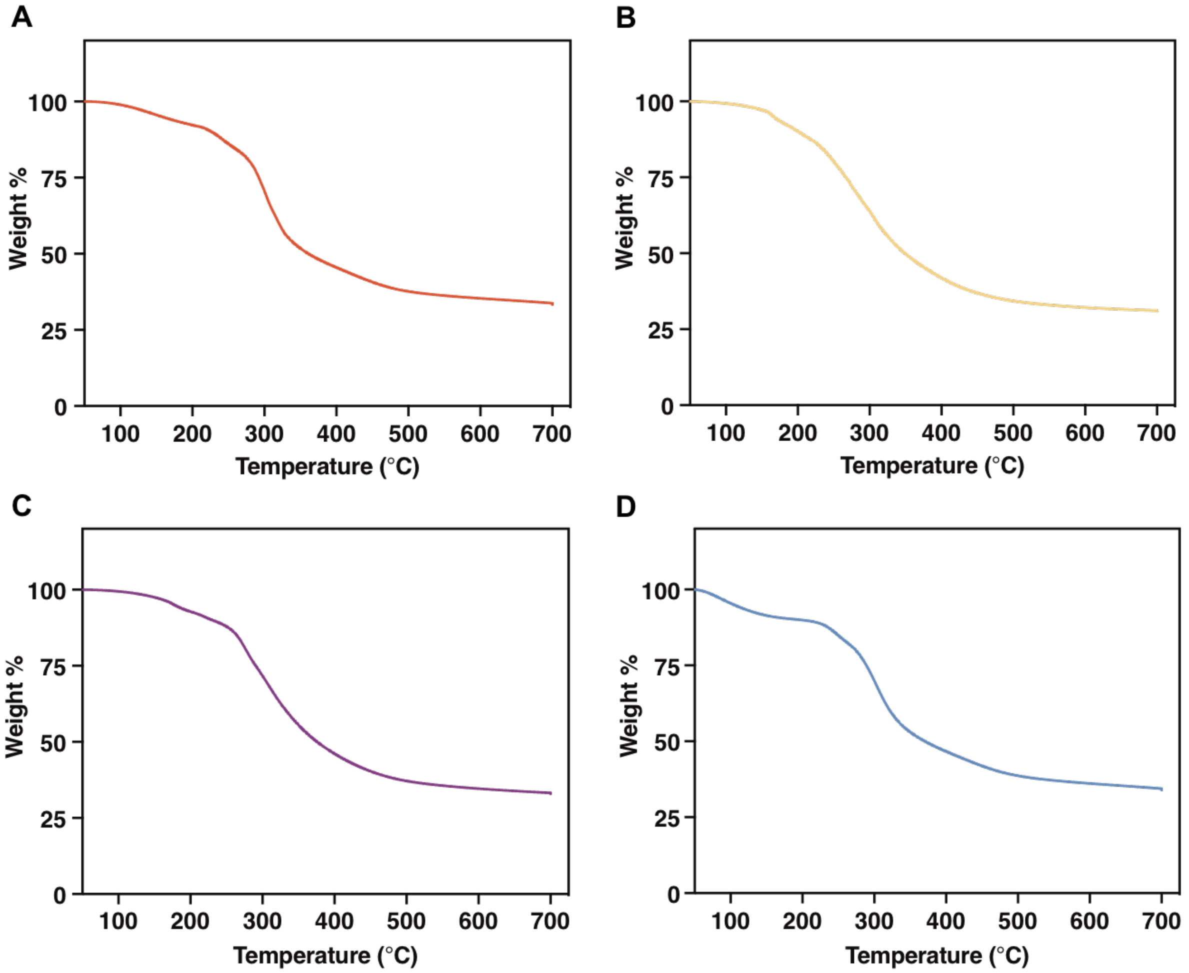


**Fig. S9. Thermal gravimetric analysis (TGA) of SLMs.** TGA of (A) EC-SLM, (B) LR-SLM, (C) SC-SLM and (D) SLM obtained from 70% ethanol treated *E. coli* cells.

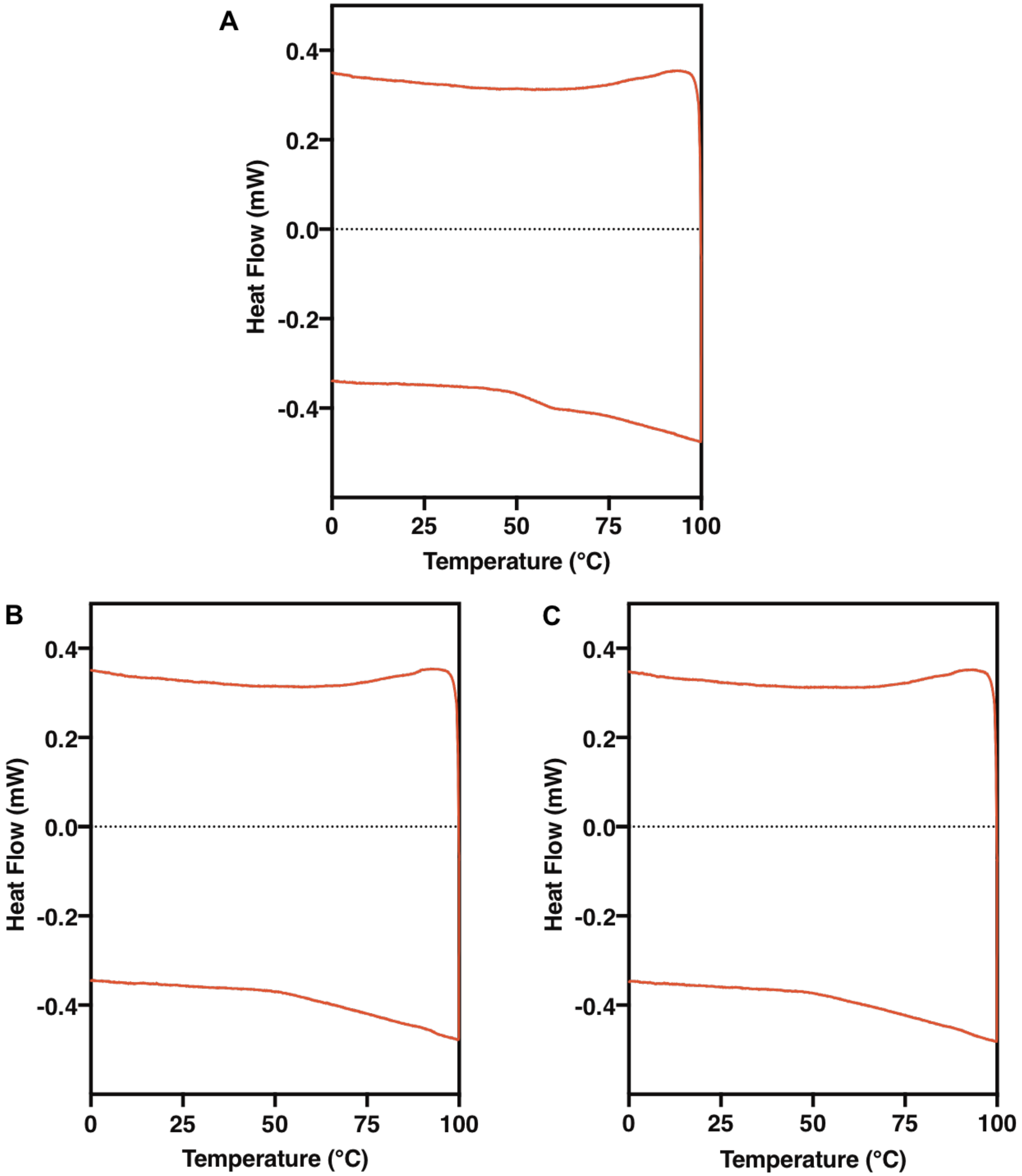


**Fig. S10. Differential scanning calorimetry (DSC) analysis of EC-SLM.** DSC curve showing heating and cooling profiles of EC-SLM (A) first cycle, (B) second cycle and (C) third cycle.

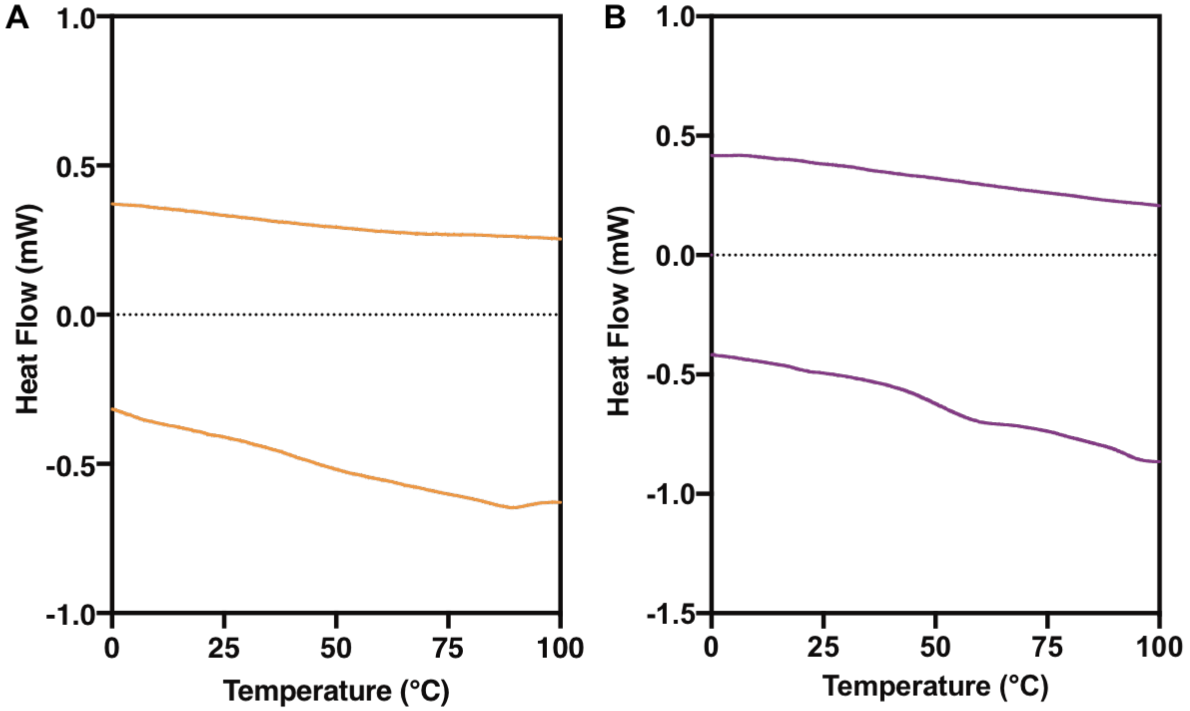


**Fig. S11. Differential scanning calorimetry (DSC) analysis of SLM.** DSC curve showing heating and cooling profiles of (A) LR-SLM and (B) SC-SLM.

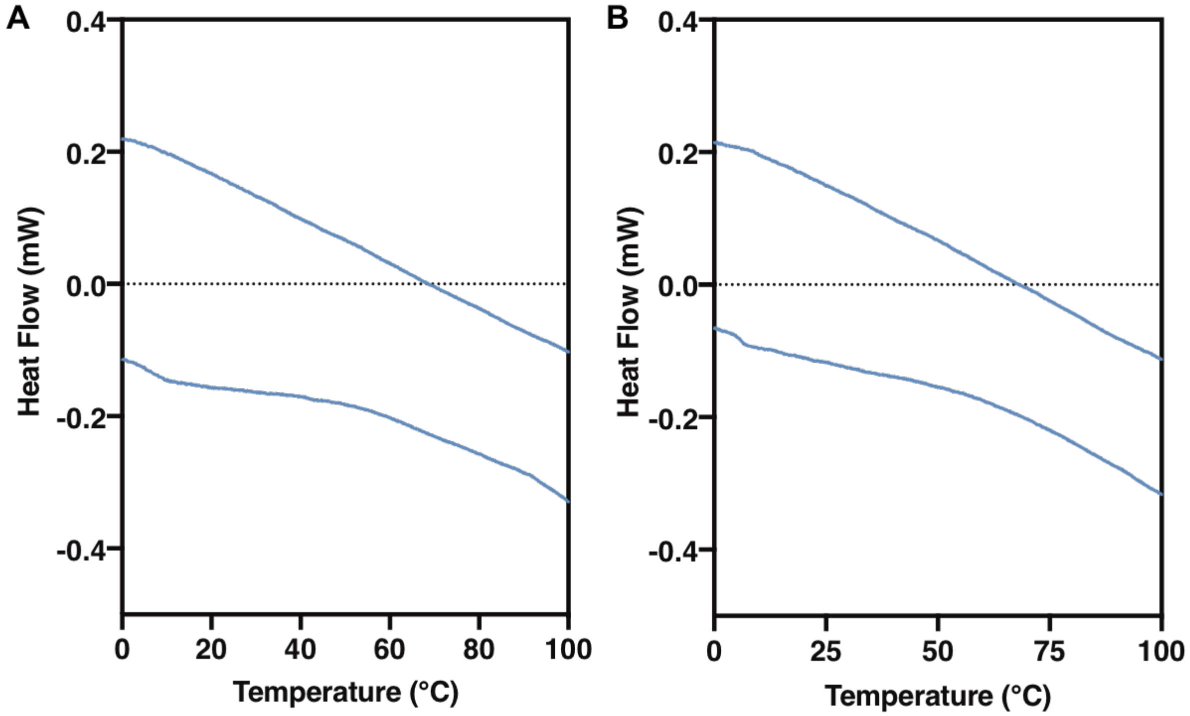


**Fig. S12. Differential scanning calorimetry (DSC) analysis of SLM obtained from 70% ethanol treated *E. coli*.** DSC curve showing heating and cooling profiles of (A) first cycle and (B) second cycle.

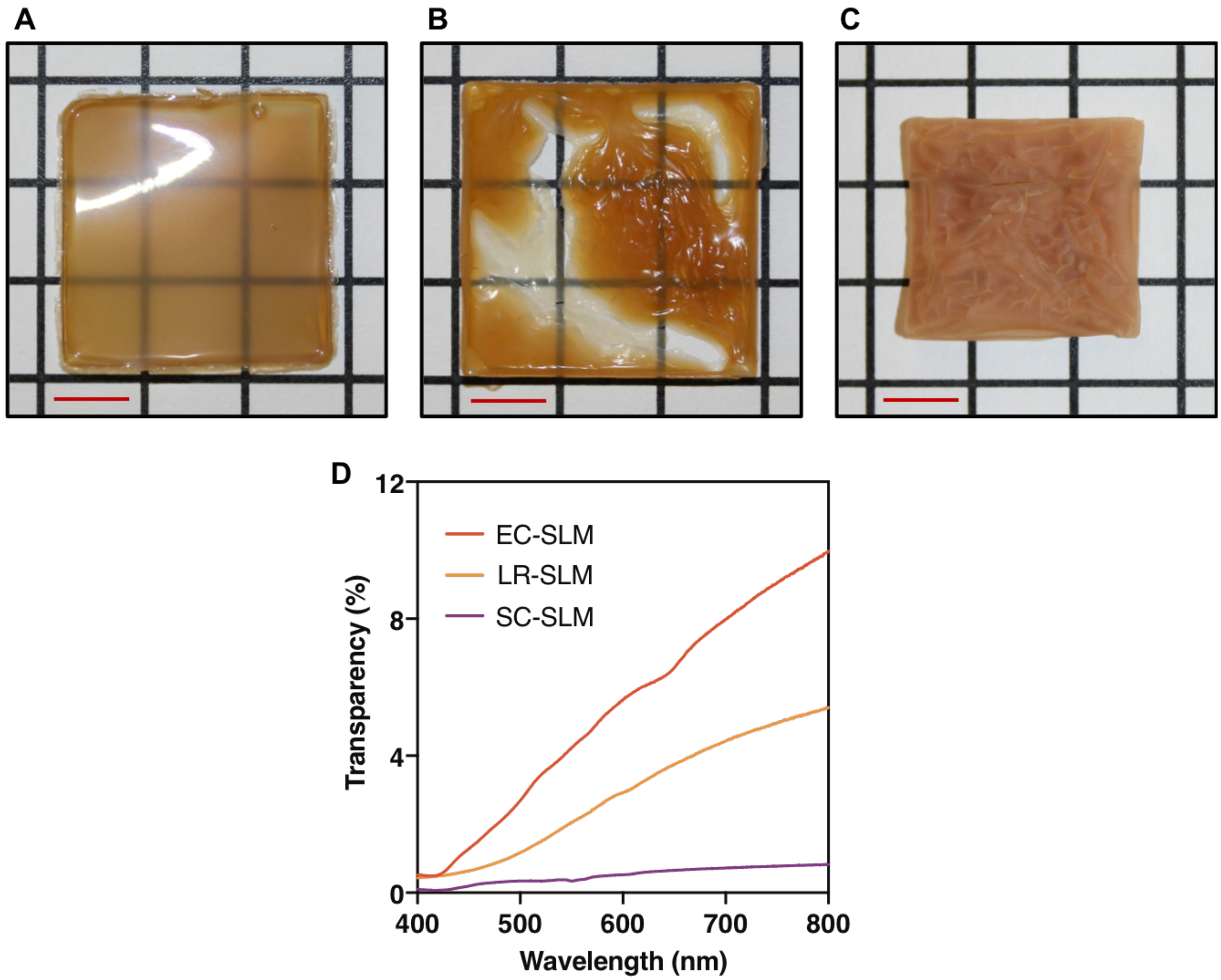


**Fig. S13. Optical transparency studies of SLMs.** Optical images show the transparency of (A) EC-SLM, (B) LR-SLM and (C) SC-SLM. (D) Absorption spectrum shows the percentage transparency of EC-SLM, LR-SLM and SC-SLM in the visible range.

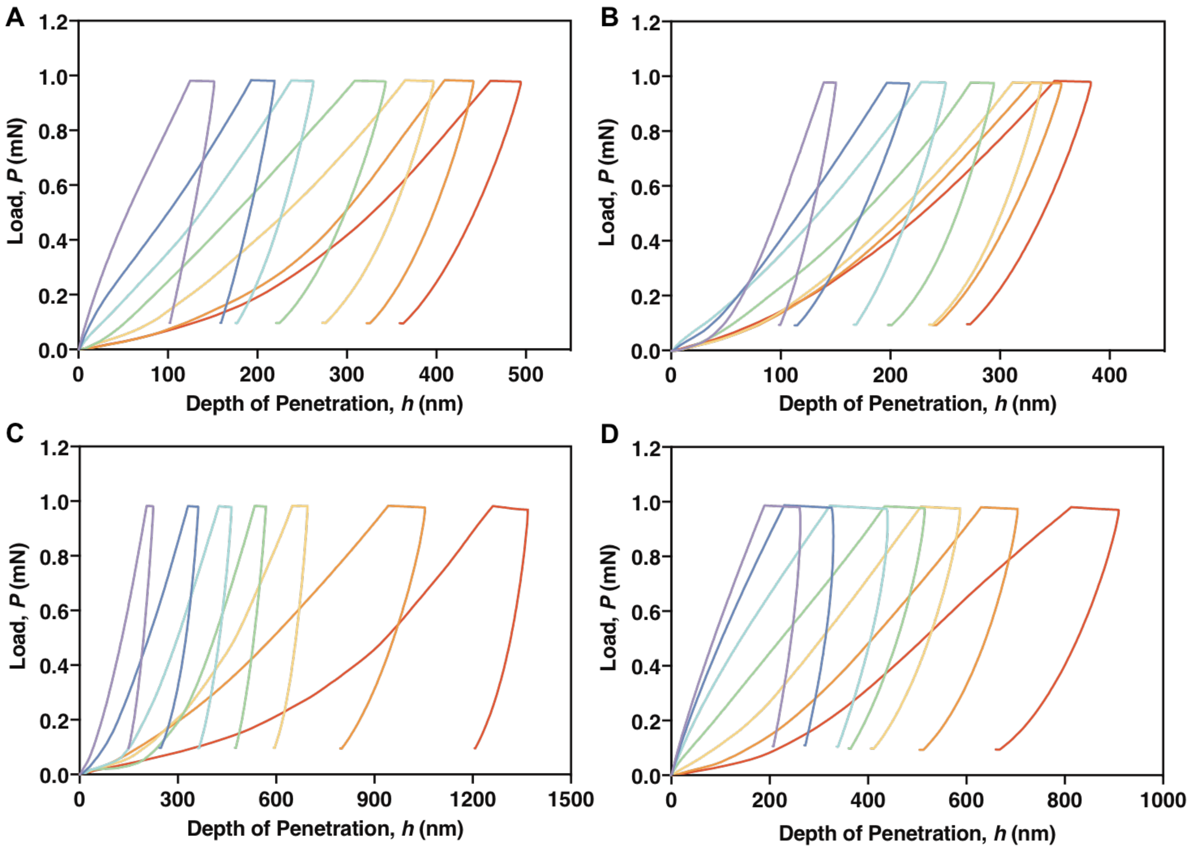


**Fig. S14. Nanoindentation studies of SLMs.** Representative load, *P*, verses depth of penetration, *h*, plots of (A) EC-SLM, (B) LR-SLM, (C) SC-SLM and (D) SLM obtained from 70% ethanol treated *E. coli*.

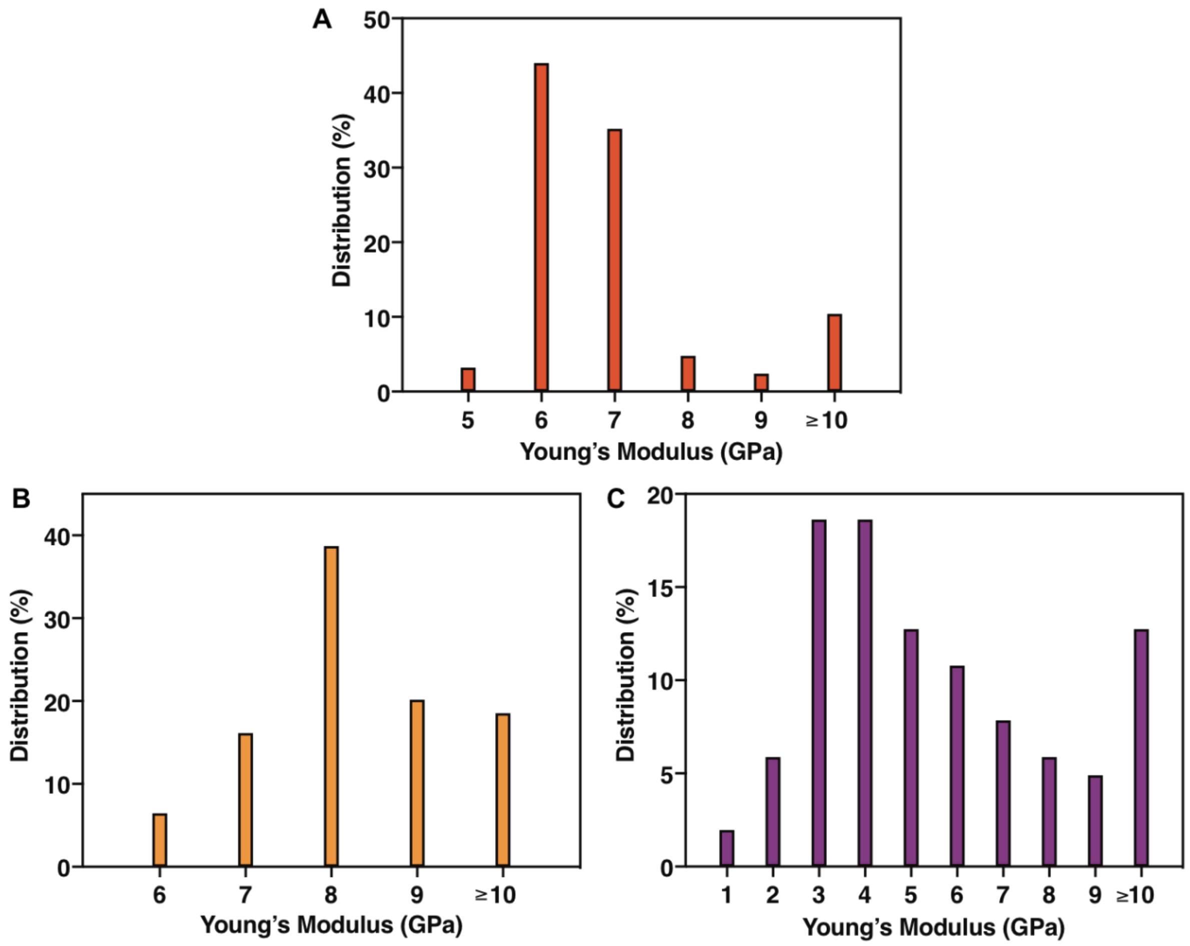


**Fig. S15. Distribution of Young's modulus of SLMs.** Binned percentage distribution of Young's modulus (data shown in Fig. 2B) obtained by nanoindentation on (A) EC-SLM, (B) LR-SLM and (C) SC-SLM. Herein, the Young's modulus value was counted as "n" for any values within n±0.5, while that above 10 GPa were also included with 10±0.5.

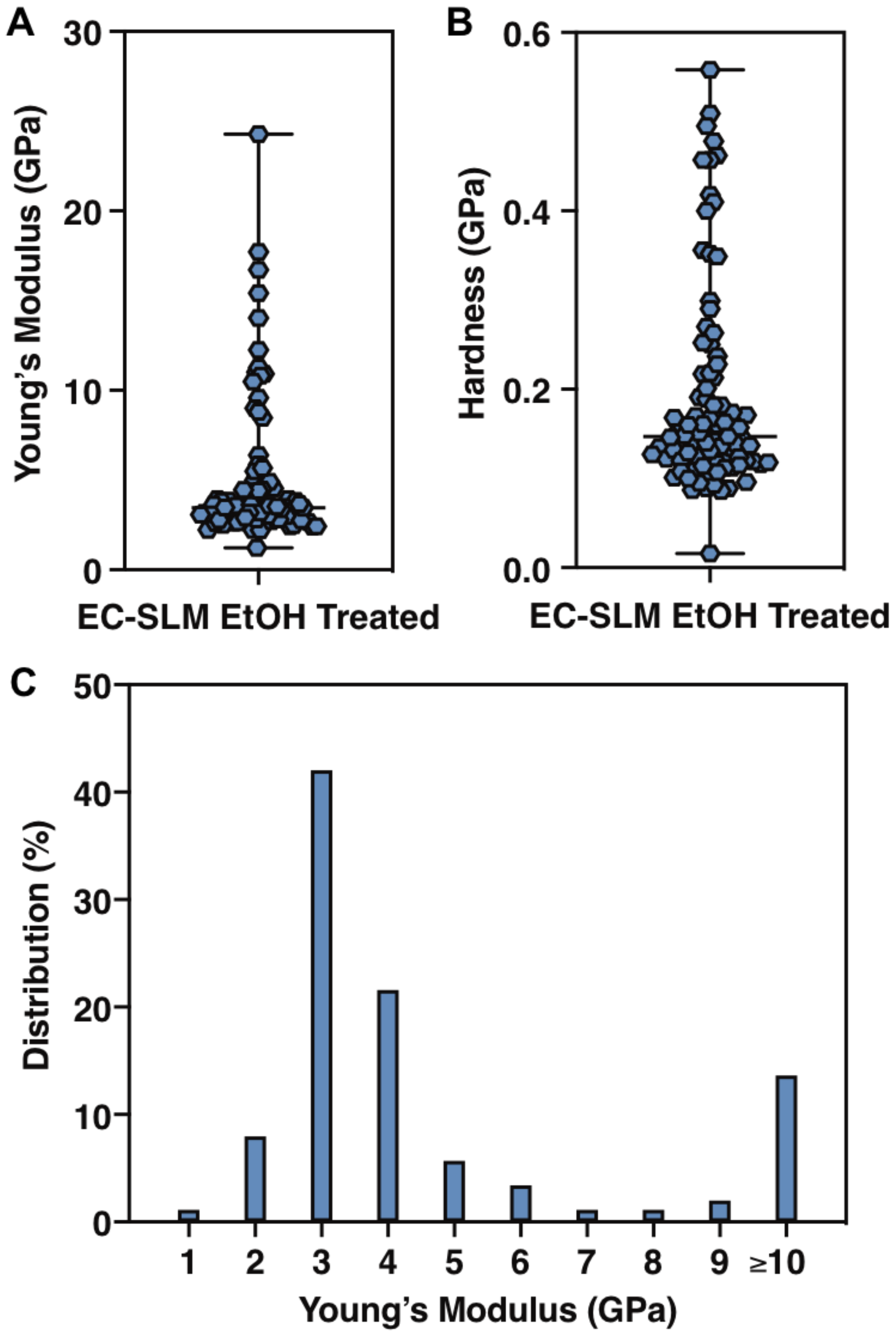


**Fig. S16. Nanoindentation analysis of SLM obtained from 70% ethanol treated *E. coli*.** (A) Young's modulus, (B) Hardness and (C) Percentage distribution of Young's modulus shown in (A). The graphs show median and the range. Herein, the Young's modulus value was counted as "n" for any values within n±0.5, while that above 10 GPa were also included with 10±0.5.

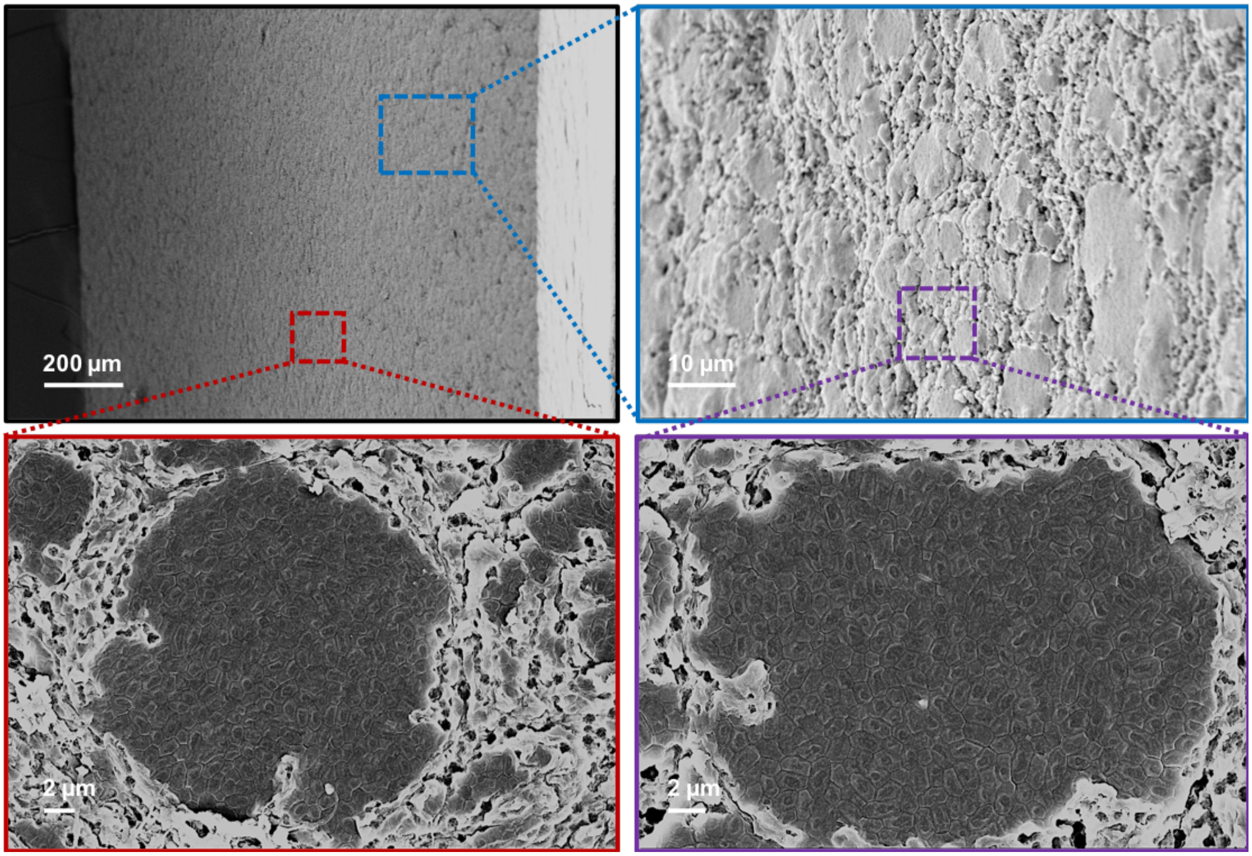


**Fig. S17. Field emission scanning electron microscopy (FESEM) images show the cross-sectional view of EC-SLM.**

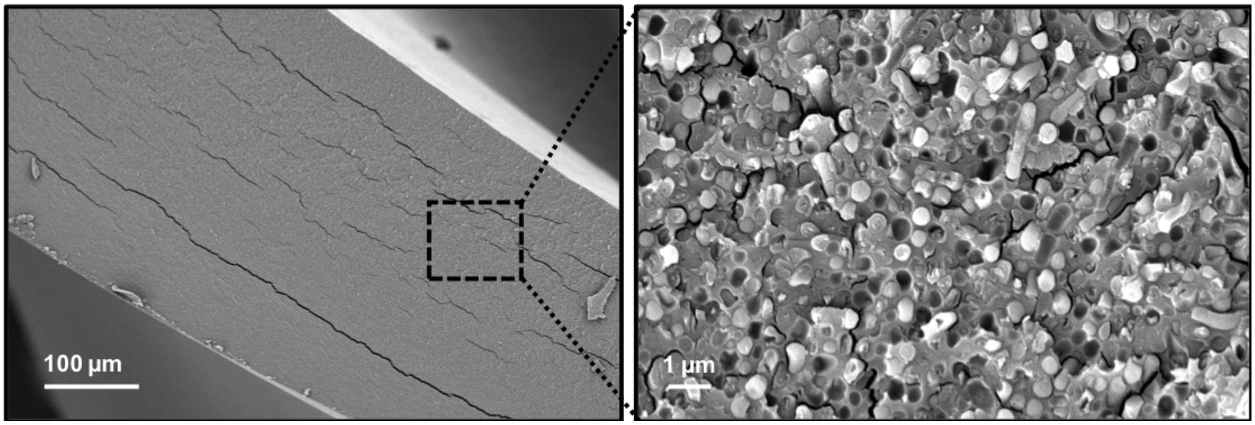


**Fig. S18. Field emission scanning electron microscopy (FESEM) images show the cross-sectional view of LR-SLM.**

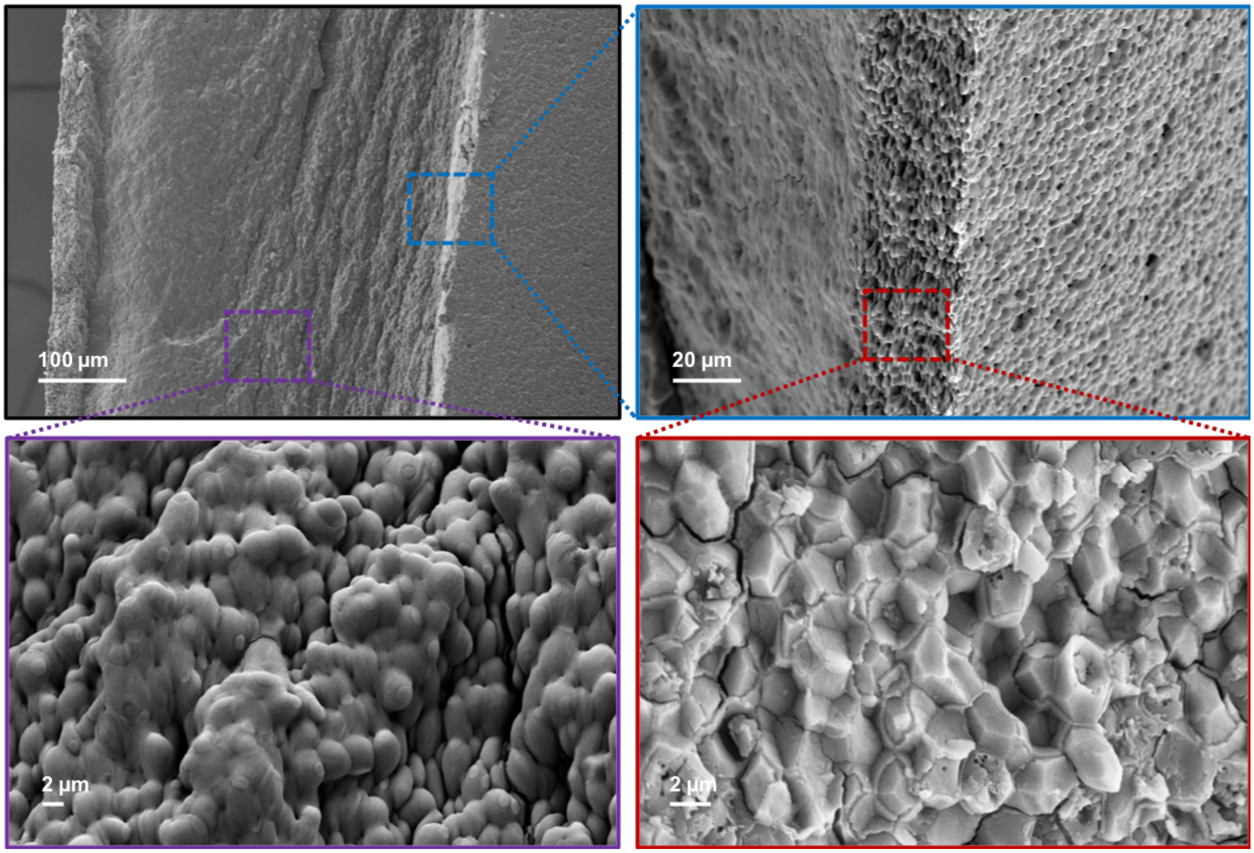


**Fig. S19. Field emission scanning electron microscopy (FESEM) images show the cross-sectional view of SC-SLM.** The outer layers (both the top and bottom) of the SLM were tightly packed, while the core was relatively less tightly packed.

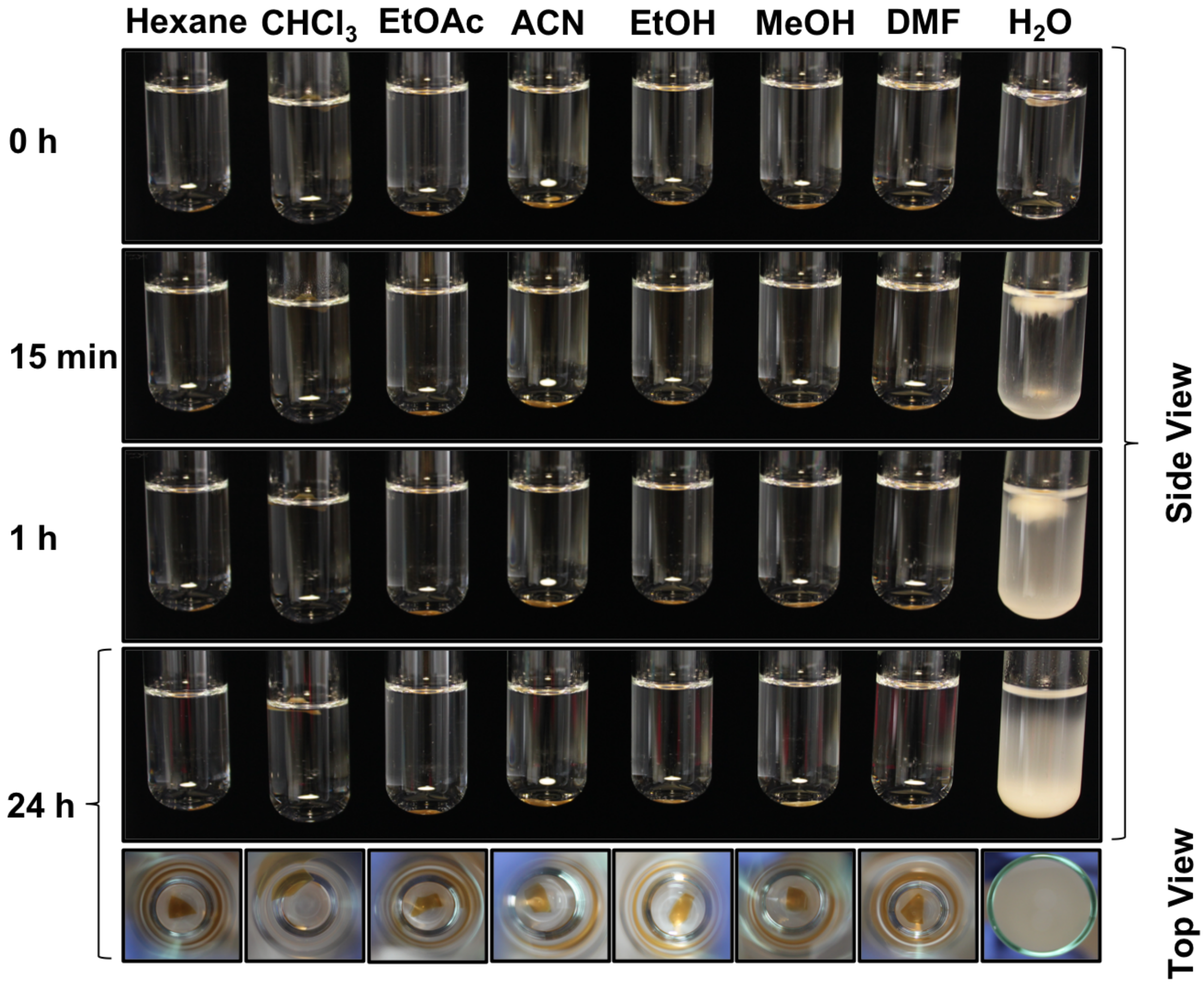


**Fig. S20. Optical images of EC-SLM in various solvents over 24 h.** $CHCl_3$: chloroform, EtOAc: ethyl acetate, ACN: acetonitrile, EtOH: absolute ethanol, MeOH: methanol, DMF: dimethylformamide.

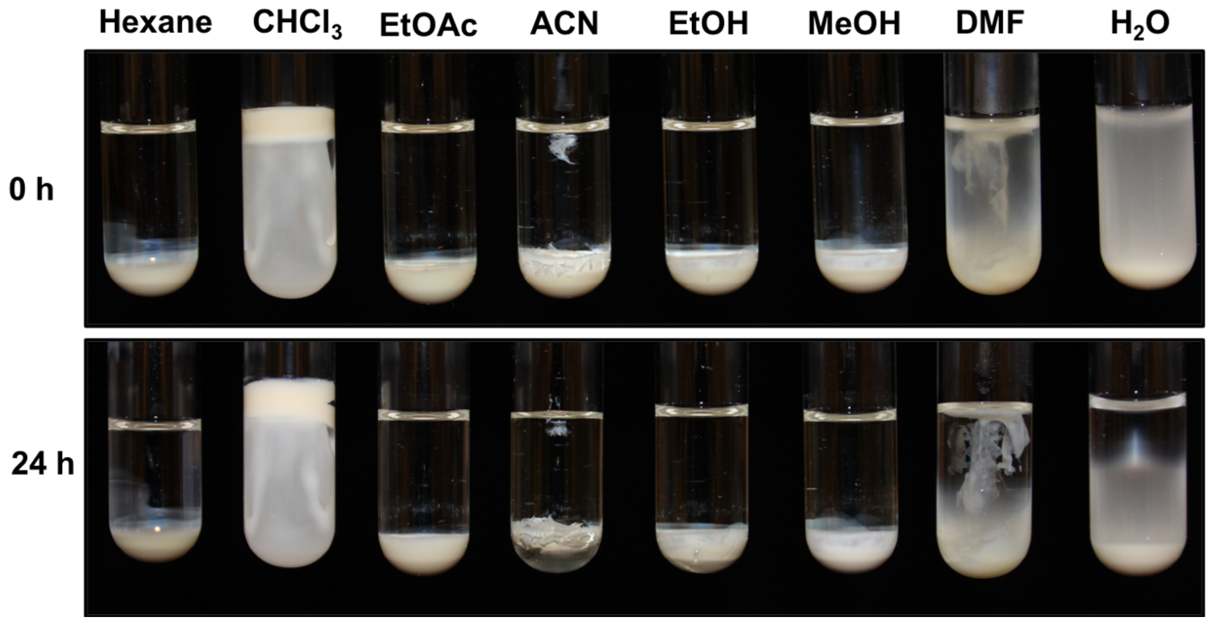


**Fig. S21. Optical images of *E. coli* pellet in various solvents.** $CHCl_3$: chloroform, EtOAc: ethyl acetate, ACN: acetonitrile, EtOH: absolute ethanol, MeOH: methanol, DMF: dimethylformamide.

| Component | Dry Weight (%) |
|---|---|
| Protein | 55 |
| RNA | 20 |
| DNA | 3 |
| Lipid | 9 |
| Lipopolysaccharide | 3 |
| Peptidoglycan | 3 |
| Glycogen | 3 |
| Metabolites | 3 |
| Others | 1 |

**Table S2. Percentage dry weight of various components in *E. coli* cell. Adapted from reference 35.**
